# Chemical Probes in Scientific Literature: Expanding and Validating Target-Disease Evidence

**DOI:** 10.64898/2026.02.19.706919

**Authors:** Melissa F. Adasme, David Ochoa, Irene Lopez, Hoang-My-Anh Do, Ellen M. McDonagh, Noel M. O’Boyle, Andrew R. Leach, Barbara Zdrazil

## Abstract

Chemical probes are indispensable tools for validating therapeutic hypotheses, yet their broader impact on early-stage drug discovery remains unquantified. To our knowledge, this study represents the first systematic, large-scale investigation of the chemical probe literature. By screening over 18 million articles using a high-quality dictionary of 561 chemical probes, we identified 20,000 articles mentioning a chemical probe which resulted in 5,558 unique target-disease (T-D) associations. Our analysis yields four principal findings that redefine the utility of these chemicals: First, we show that chemical probe evidence typically precedes the appearance of structured data in major knowledge bases by 1–7 years, providing a crucial lead time for target prioritisation. Second, we identified 353 T-D pairs (6.4%) with no prior evidence in the Open Targets Platform, highlighting the approach’s discovery potential. Third, the application of strict novelty filters uncovered 135 new high-confidence associations between targets and diseases, revealing distinct opportunities for therapeutic repurposing in non-oncological, rare autoimmune diseases, and diseases without effective therapies due to complex biology or high treatment resistance. Finally, we demonstrate that chemical probes are essential for strengthening evidence, providing functional validation for associations previously supported only by weaker, correlative data such as RNA expression or animal models. Collectively, these findings illustrate that chemical probes catalyse early therapeutic discovery, emphasising the importance of cataloguing existing probes and identifying new ones.

## Introduction

Chemical probes - selective, potent, and cell-permeable small molecules - are indispensable tools in modern biological research and drug discovery.^1–4^ They empower researchers to dissect complex biological systems by specifically modulating target protein function, offering advantages like temporal control and dose-dependent effects that complement traditional genetic techniques. Their applications are broad, critically including target validation, pathway mapping, and the investigation of disease mechanisms, making them central to establishing and exploring links between molecular targets and pathological states.

The effective use of these powerful molecular tools hinges on rigorous characterization and an awareness of potential limitations. Key "fitness" parameters, including target potency, selectivity over related proteins, and cell permeability, must be well-defined. Critically, the potential for off-target interactions necessitates careful validation, often involving specific negative controls like inactive structural analogues, to ensure any observed effects are truly linked to the intended target.^4–6^ Global public-private partnerships such as the Structural Genomics Consortium (SGC) (https://www.thesgc.org),^7^ as well as public resources such as the Chemical Probes Portal (chemicalprobes.org)^1^ and Probes & Drugs (https://www.probes-drugs.org)^3,8^ provide valuable guidance by curating and recommending well-characterized chemical probes, promoting transparency and facilitating robust experimental design.^9^ Reporting the characteristics of chemical probes transparently is crucial for their reliable use by the wider scientific community.

However, despite the establishment of these foundational principles, a significant challenge arises when attempting to assess the *cumulative* evidence and real-world impact derived from the widespread application of chemical probes, as documented across the vast and exponentially growing body of scientific literature. Researchers routinely report findings using established chemical probes in diverse experimental settings, often providing crucial data points that incrementally build the case for novel target-disease (T-D) links, validate (or invalidate) therapeutic hypotheses, or elucidate fundamental biological pathways. The confidence in such findings, particularly in the context of target validation, grows substantially when multiple, structurally distinct chemical probes targeting the same protein yield consistent biological outcomes across various experimental models and laboratories. However, this wealth of validation data remains largely dispersed and buried within individual publications, hindering systematic analysis and may diminish the wider appreciation of the true value of chemical probes in the life sciences and especially drug discovery: they enable precise exploration of biological targets and pathways before major drug development investments are made.

The systematic aggregation and interpretation of diverse lines of evidence are paramount for robust target validation and prioritisation in drug discovery. The Open Targets Platform (https://platform.opentargets.org/),^10^ established through an academic-industry partnership, integrates and harmonises a vast array of evidence types to illuminate the intricate relationships between molecular targets and human diseases/traits. This comprehensive platform curates and presents evidence from germline genetics, somatic mutations, RNA expression, animal models, approved drugs (including clinical candidates), and crucially, the scientific literature to contextualize target-disease relationships. However, the literature-derived evidence, while broad, often lacks the focused mechanistic validation provided by well-characterised chemical probes.

This study intends to address this gap by systematically exploring the published scientific literature to identify and analyse articles where chemical probes have been instrumental in testing T-D relationships. Using advancements in Natural Language Processing (NLP) techniques, which enable the large-scale extraction and analysis of information from unstructured text, we examine patterns in how these molecular tools are utilised experimentally as described in scientific literature. The value of this work lies in providing a consolidated view of chemical probe-based evidence supporting distinct T-D links. It therefore offers insights into the practical drug discovery applications of these critical reagents, highlighting robustly supported associations as well as areas of emerging focus in more recent years. The major strength of this analysis lies undoubtedly in its ability to identify these emerging T-D links, which shall guide future decisions on investments in drug discovery and development programmes.

## Results

### Pilot study exploring references to SGC chemical probes in open-access scientific literature

A pilot study was performed using a small set of 56 high-quality (HQ) chemical probes (i.e. recommended by the SGC, https://www.thesgc.org/) to gain preliminary insights about the feasibility and throughput of using NLP to extract chemical probe-target-disease (P-T-D) triples from the literature, and also to understand better the context in which chemical probes are being reported (Figure 1, left panel). Using very strict selection criteria (see Methods), we analysed a corpus of 75,141 open-access articles from EuropePMC, containing dictionary-based annotations for chemicals/drugs, genes/proteins, and diseases. This led to the identification of 108 publications mentioning at least one of the 56 SGC chemical probes (as chemical entity) alongside their corresponding target within the same sentence (Supplementary File pilot_triples.csv). A single article may contain multiple such sentences referring to the same chemical probe-target (P-T) pair, and individual articles often mention a variety of different chemical probes and their respective targets. These initial findings correspond to 142 unique publication-chemical probe-target triples derived from these 108 articles, covering 22 unique chemical probes and 24 unique targets.

**Figure 1:**
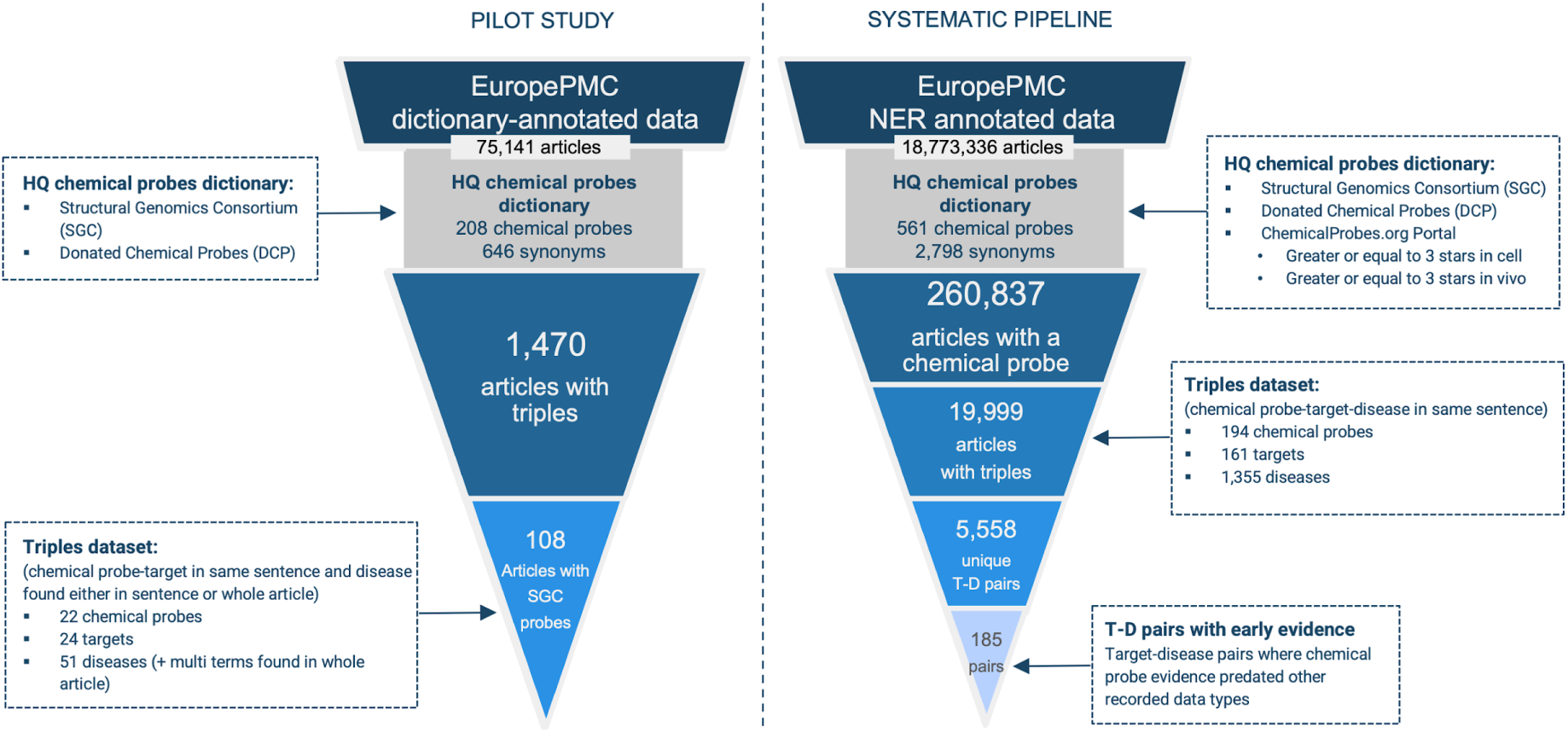
Overview of the pipeline for identifying early T-D associations from chemical probe literature. (Left) The pilot study used 75,000 dictionary-based annotated articles to extract sentences containing P-T-D information. It served as a proof-of-concept, involving the manual curation of 100 publications to evaluate the feasibility of the text-mining approach and to create a Gold Standard for validation. (Right) The systematic pipeline was applied to an NER annotated corpus of ∼ 19 million open-access publications to extract sentences containing P-T-D information, which were processed to yield 5,558 unique T-D pairs.

Investigation of the disease context revealed that 56 of the 142 unique associations also explicitly mentioned a disease within the sentence containing the chemical probe and the target. For the remaining 86 associations, although a disease was not present in that specific sentence, a relevant disease entity was annotated elsewhere within the full text of the article. Subsequent manual curation of the 108 articles indicated that 74 (69%) presented a clear, strong link between the chemical probe, its target, and the associated disease context (e.g., cases where entities appeared within the same sentence or nearby). By contrast, the remaining 31% of articles described a weaker or more indirect link, where the relationship between the entities was less explicit, buried within the full text, and more challenging to extract. Figure 2 shows two examples of the articles found: on the left, an example describing a strong and direct link, while the example on the right is representative of a weaker, indirect link.

**Figure 2.**
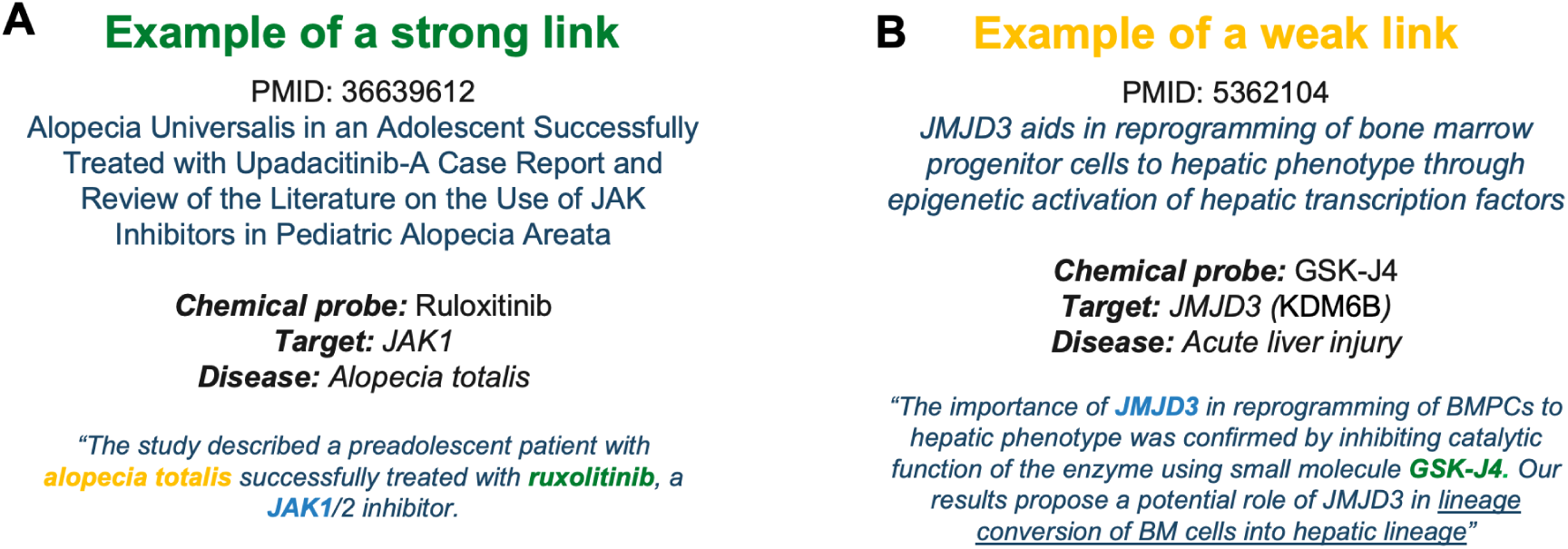
Examples of strong and weak P-T-D links identified in scientific articles. (A) Strong link: Chemical probe “ruloxitinib”, target “JAK1” and disease “alopecia totalis” are explicitly connected within a single sentence. (B) Weak/indirect link: Chemical probe “GSK-J4” and target “JMJD3 (KDM6B)” are mentioned together, with the related disease “Acute liver injury” inferred from context elsewhere in the article.

Manual curation of the test set identified 90 articles that also reported bioactivity data associated with chemical probes binding to their known primary targets, off-targets, and other reported targets. These 90 articles correspond to 71 unique PubMedID-chemical probe-target triples, varying in the context of diseases from which specific bioactivity information could be successfully extracted. For 16 articles processed from the test set, however, associated bioactivity data could not be retrieved. The analysis identified several reasons for the lack of extractable bioactivity data. In some cases, the chemical probe was only mentioned peripherally and was not part of the core analysis. In other instances, the chemical probe was used, but bioactivity experiments were performed on different targets. Finally, some articles discussed the chemical probe’s use but did not provide explicit, identifiable bioactivity values.

### Systematic exploration of probe-target-disease links in the scientific literature

Building upon the insights gained from the pilot study regarding P-T-D co-mentions, we expanded the scope of our investigation to systematically analyse a significantly larger literature corpus of open-access biomedical literature using automated methods (Figure 1, right panel). To scale up, we leveraged the Open Targets literature-mining corpus (18,773,336 articles), which includes a set of entities tagged using Named Entity Recognition (NER), namely chemicals/drugs, genes/proteins, and diseases.^11^ An expanded and curated High Quality (HQ) chemical probe dictionary served as a key tool for the initial processing step (Supplementary File “probes_HQ.csv”). This dictionary, compiled from established resources, includes 89 SGC chemical probes, 110 Open Science chemical probes, and 508 chemical probes extracted from the ChemicalProbes Portal, which accounted for 561 unique chemical probes (having distinct InChIKey’s), 554 primary names and 2,798 associated synonyms. Perhaps surprisingly, the three sources have only 5 chemical probes in common, with 18 chemical probes being present solely in the SGC set, 9 solely in the Open Science donated chemical probes set, and 363 solely in the ChemicalProbes Portal (Supplementary Figure S1).

Using this dictionary as a filter for the NER-annotated articles enabled the identification of 260,837 articles containing mentions of known HQ chemical probes or their synonyms. Within these publications, a total of 1,636,358 chemical probe mentions were detected, reflecting the common occurrence of multiple mentions per article; these mentions corresponded to 194 unique chemical probe entities. This initial filtering created a focused subset of relevant articles, thereby streamlining the subsequent computationally intensive search for P-T-D relationships.

The second analysis stage was focused on the extraction of P-T-D triples from this subset and identified potential associations within 103,381 articles, encompassing 5,709 unique chemicals (not only chemical probes), 9,238 unique targets, and 3,574 unique diseases (Supplementary File “ner_all_triplets.tsv”). The HQ Chemical Probe Dictionary was used for an additional filtering step to enhance the confidence of these findings, by retaining only those chemical probe-target associations where the mentioned target is also reported as the main target of the specific chemical probe. This stringent curation resulted in a final high-confidence NER chemical probes dataset, derived from 19,999 articles (12,249 primary articles and 7,750 review articles), that links 194 unique chemical probes to 161 validated targets and 1,353 distinct diseases (Supplementary File “ner_probes_triplets_ptpairs.tsv”).

Further analysis characterised the distribution patterns of the identified chemical probes within the open biomedical literature corpus and their associated biological entities (Figure 3). The frequency with which individual chemical probes have been mentioned across different articles is depicted in Figure 3A.

**Figure 3.**
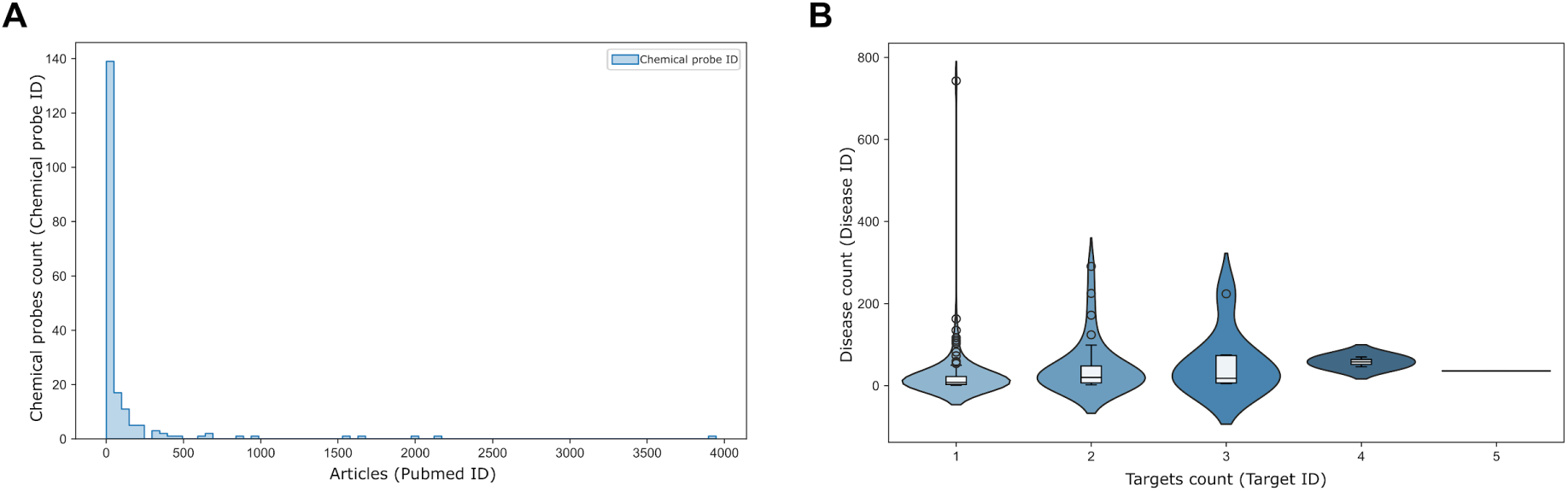
Distribution analysis of identified chemical probes and their associated targets and diseases. (A) The histogram shows the distribution of chemical probes based on the number of articles in which they were identified (range: 1 to 3,946 articles per chemical probe). (B) The violin plot shows the number of linked diseases per chemical probe, grouped by the number of unique targets (range: 1 to 5) and unique diseases (range: 1 to 744) associated with them. Most chemical probes are associated with a few targets (1 to 3) and a relatively low number of diseases (median=10) in the dataset.

While the number of associated articles per unique chemical probe varied widely, ranging from a minimum of 1 to a maximum of 3,946 publications (median=16.5), the density distribution is highly skewed.

This indicates that the vast majority of identified chemical probes appear in only a small number of articles, with relatively few chemical probes being mentioned frequently across the literature (all chemical probes frequency in Supplementary File “ner_probes.tsv”). Complementing this, the relationships between chemical probes and their linked targets and diseases were examined (Figure 3B). These violin plots illustrate that chemical probes are typically associated with only a small range of targets (1 to 5), an observation consistent with the definition of chemical probes as selective tool compounds.

The number of associated diseases per chemical probe showed greater variability, ranging from 1 to 744. Despite this range in disease associations, the violin plot reveals a strong concentration in the region corresponding to low target counts (1-3) and low disease counts (median=10). This indicates that most chemical probes within this dataset are linked to a single target and a limited number of diseases, suggesting focused investigation contexts in the analysed scientific literature .

Analysing the frequency of the occurrence of sentences reporting relevant P-T-D links in scientific articles (Figure 4), the "Other" category yielded the highest frequency (30.2%). This category comprises text from articles that do not follow a conventional IMRaD (Introduction, Methods, Results, and Discussion) structure. This finding indicates that a significant portion of evidence is located within publications where results are presented within topically titled subsections rather than within a single, overarching "Results" section. Such a format is frequently employed by journals like *Frontiers in Oncology*, *World Journal of Gastrointestinal Oncology*, and *Drug Design, Development and Therapy*, among many others. For traditionally structured publications, significant concentrations of evidence sentences were identified in key narrative sections: "Discussion" (17.8%), "Introduction" (16.5%), and "Abstract" (14%). Substantial counts were also registered for the "Results" (10%) and "Methods" (4.9%) sections, reflecting their roles in reporting experimental findings and procedures related to the investigated chemical probe(s). Lower, yet notable, frequencies characterised the "Conclusion" (2.7%) and "Title" (2.1%) sections. In contrast, sections primarily associated with data presentation adjuncts ("Figures" legends, "Tables" content) or article metadata ("Abbreviations", "Case study", "Supplementary" information, "Acknowledgments", "Author contribution") yielded minimal counts of the targeted evidence sentences, typically fewer than 250 instances per section.

**Figure 4.**
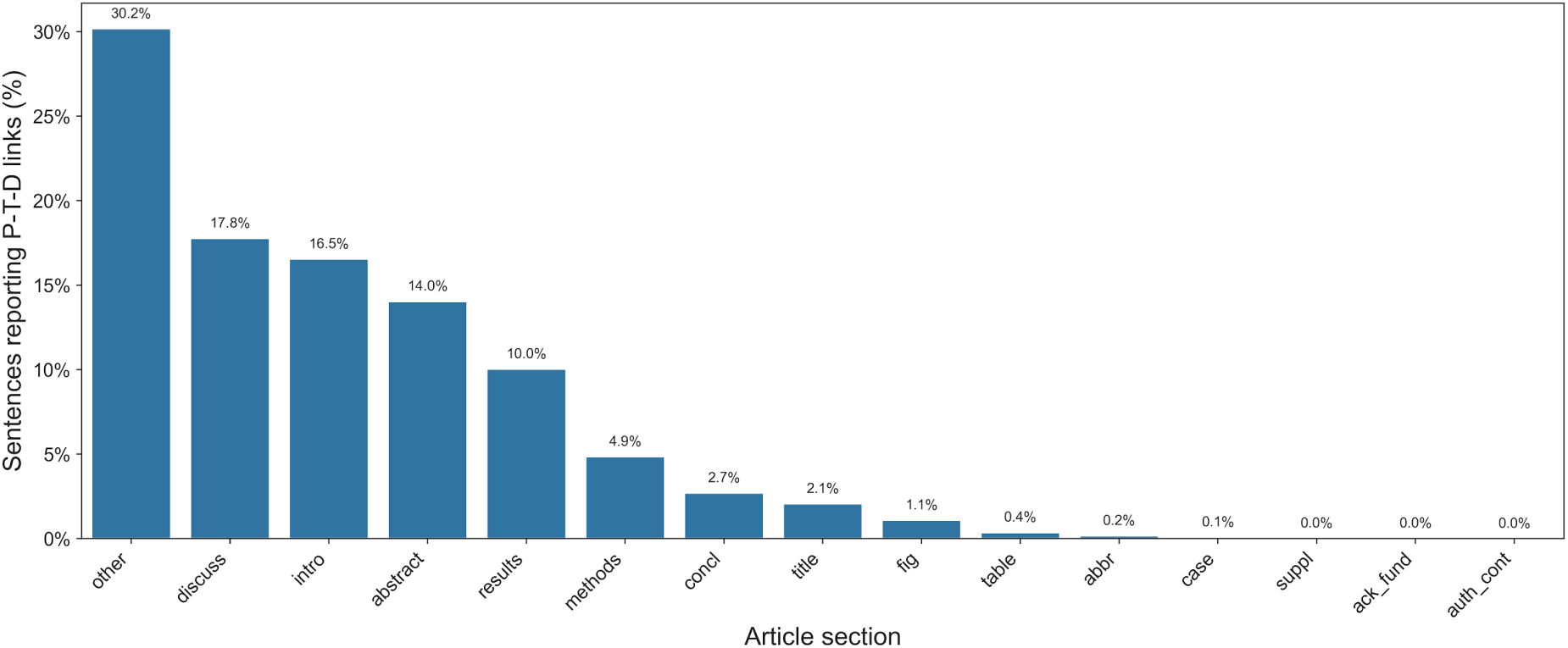
Distribution of sentences within scientific articles containing evidence for T-D links mediated by chemical probes. Sentences are grouped and counted according to the article section from which they originated.

### Uncovering target-disease links with published chemical probes

Analysis of published chemical probe experiments enables the identification of functional links between target modulation and disease phenotypes, providing opportunities to discover novel associations or strengthen existing ones. Our dataset comprised 5,558 unique T-D pairs discussed in the context of chemical probe investigations across the analysed literature. To facilitate readability and standardisation, throughout this manuscript targets are referred to by their HGNC gene symbols,^12^ diseases by their Experimental Factor Ontology (EFO) terms,^13^ and chemical probes by the ChEMBL compound preferred name.^14^

Among the 161 proteins investigated, a distinct set of targets emerged as the most frequently studied using chemical probes (Supplementary File ner_targets.tsv). This list is dominated by well-established key players in critical cellular pathways, including protein kinases central to signalling and cell proliferation, such as members of the Janus kinase family (JAK1, JAK2, JAK3), the epidermal growth factor receptor (EGFR), and anaplastic lymphoma kinase (ALK). Other prominent targets include key regulators of cell growth and metabolism (MTOR, PIK3CA), the cell cycle (CDK4), apoptosis (BCL2), and epigenetic mechanisms (EZH2). The frequent appearance of these proteins underscores the research community’s focus on modulating key targets in cancer and inflammatory disease pathways.

Correspondingly, the analysis of 1,353 disease phenotypes revealed a strong focus on oncology (Supplementary File “ner_diseases.tsv”). The most frequently mentioned diseases included general terms such as "neoplasms" and "cancer", as well as specific malignancies, including "non-small cell lung carcinoma", "acute myeloid leukaemia", "chronic lymphocytic leukaemia", "breast cancer", "hepatocellular carcinoma", and "triple-negative breast cancer". This prevalence of cancer-related research aligns with the identified target landscape, which is rich in proteins implicated in oncogenesis. Notably, inflammatory and autoimmune conditions are also represented, with Rheumatoid Arthritis and the haematological disorder Myelofibrosis featuring prominently. The co-occurrence of these diseases with targets like the JAK kinases highlights the utility of chemical probes in exploring therapeutic hypotheses that span multiple disease areas.

The amount of supporting evidence varied significantly, with the number of articles mentioning a specific T-D pair in conjunction with a chemical probe ranging from 1 to 1,260. The 20 most frequent T-D pairs are listed in Table 1 (full list in Supplementary File “ner_target_disease_pairs.tsv”), which highlights that a single T-D relationship is often investigated using multiple distinct chemical probes across numerous publications, reflecting the availability of different tool compounds for targets and the diversity of experimental approaches. Table 1 details the specific chemical probes identified for each example pair, reporting the highest clinical phase registered (if available) for the chemical probe in any indication (value in parenthesis) and the count of unique sentences where the relevant P-T-D triple context was found within the cited articles (value in square brackets). Therefore, the information under "Chemical probe (max clinical phase)[sentences]" indicates the breadth of chemical probe-based evidence, serving as a proxy for the strength of the link. In parallel, the "Article Count" reflects the overall volume of research, which helps gauge the maturity or novelty of the investigation when publication dates are considered.

**Table 1.**
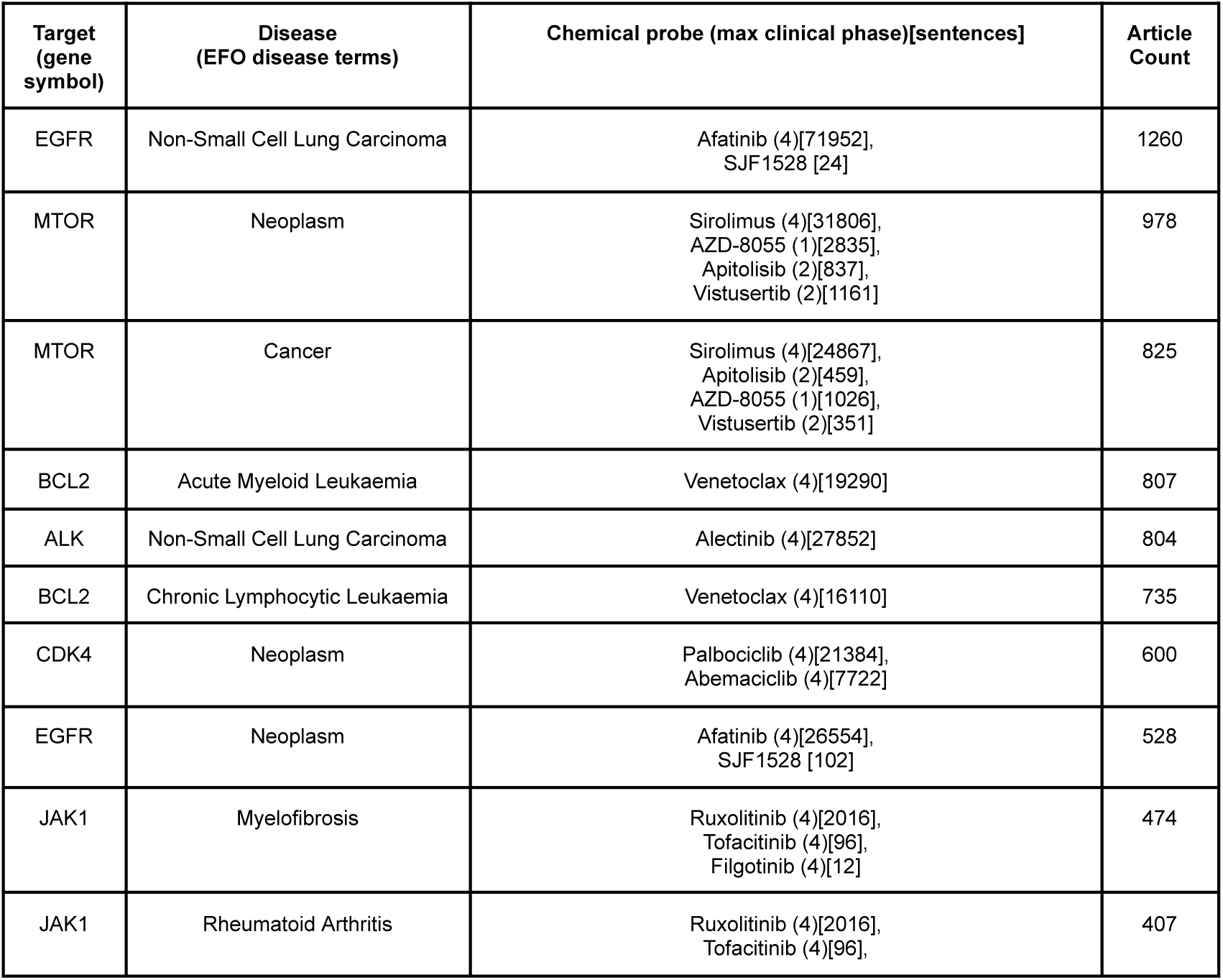
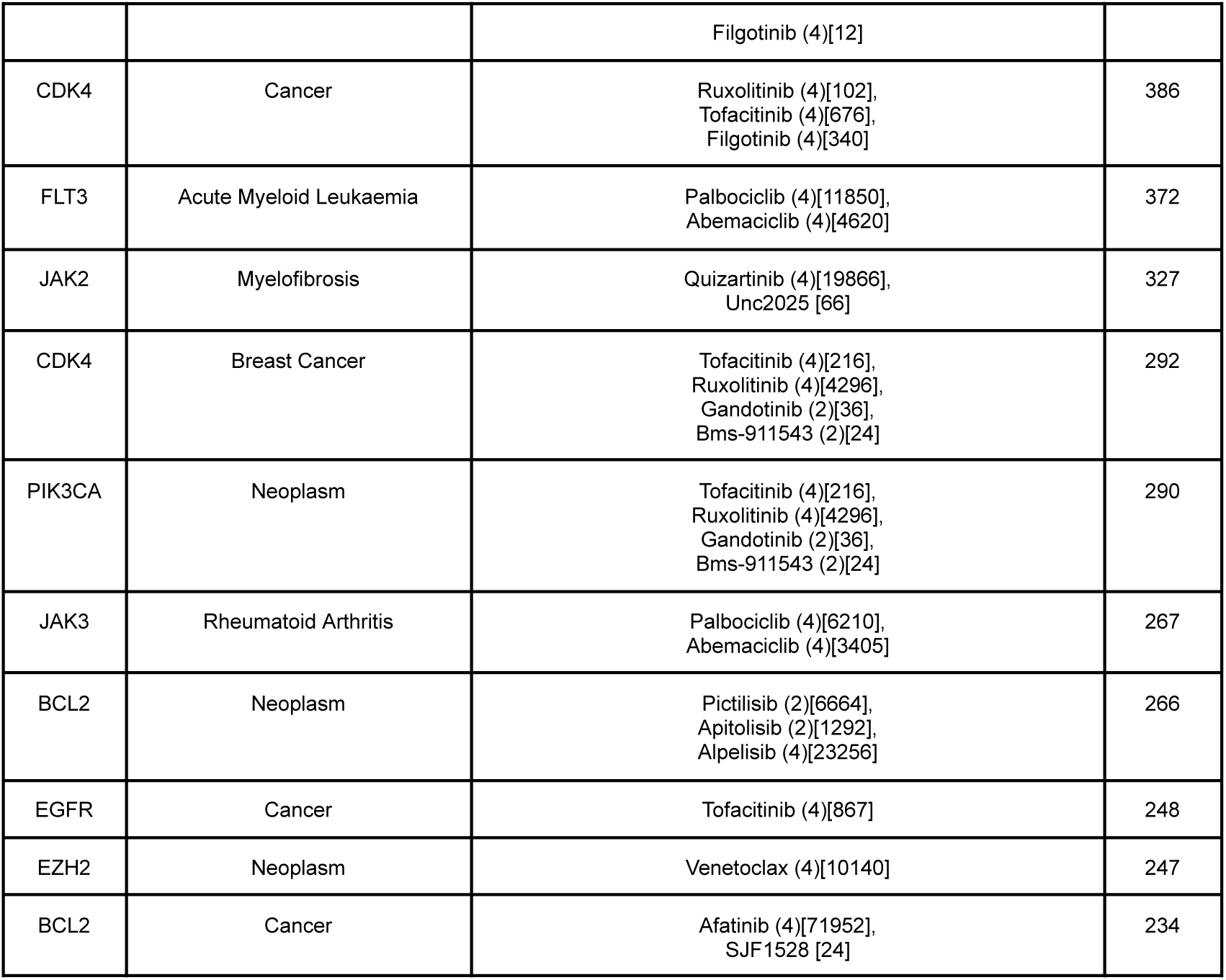
The 20 most frequent T-D pairs with associated chemical probes, the maximum clinical phase of the chemical probe and the corresponding number of articles these pairs appeared in. The highest clinical phase reached for a respective compound in clinical investigations (across all indications) is indicated in parentheses (1…phase 1; 2…phase 2; 3…phase 3; 4…approved). The number of unique sentences in which the chemical probe was mentioned together with the T-D pair can be found in square brackets.

**Table 2.**
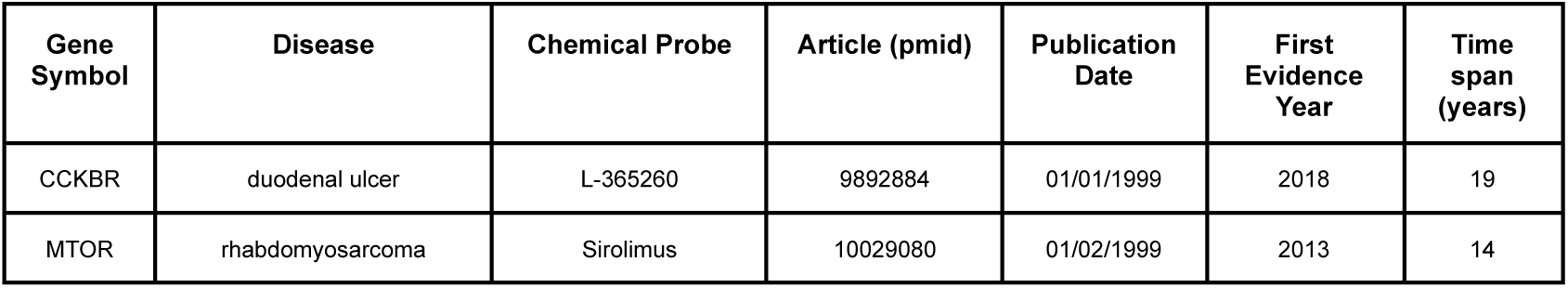
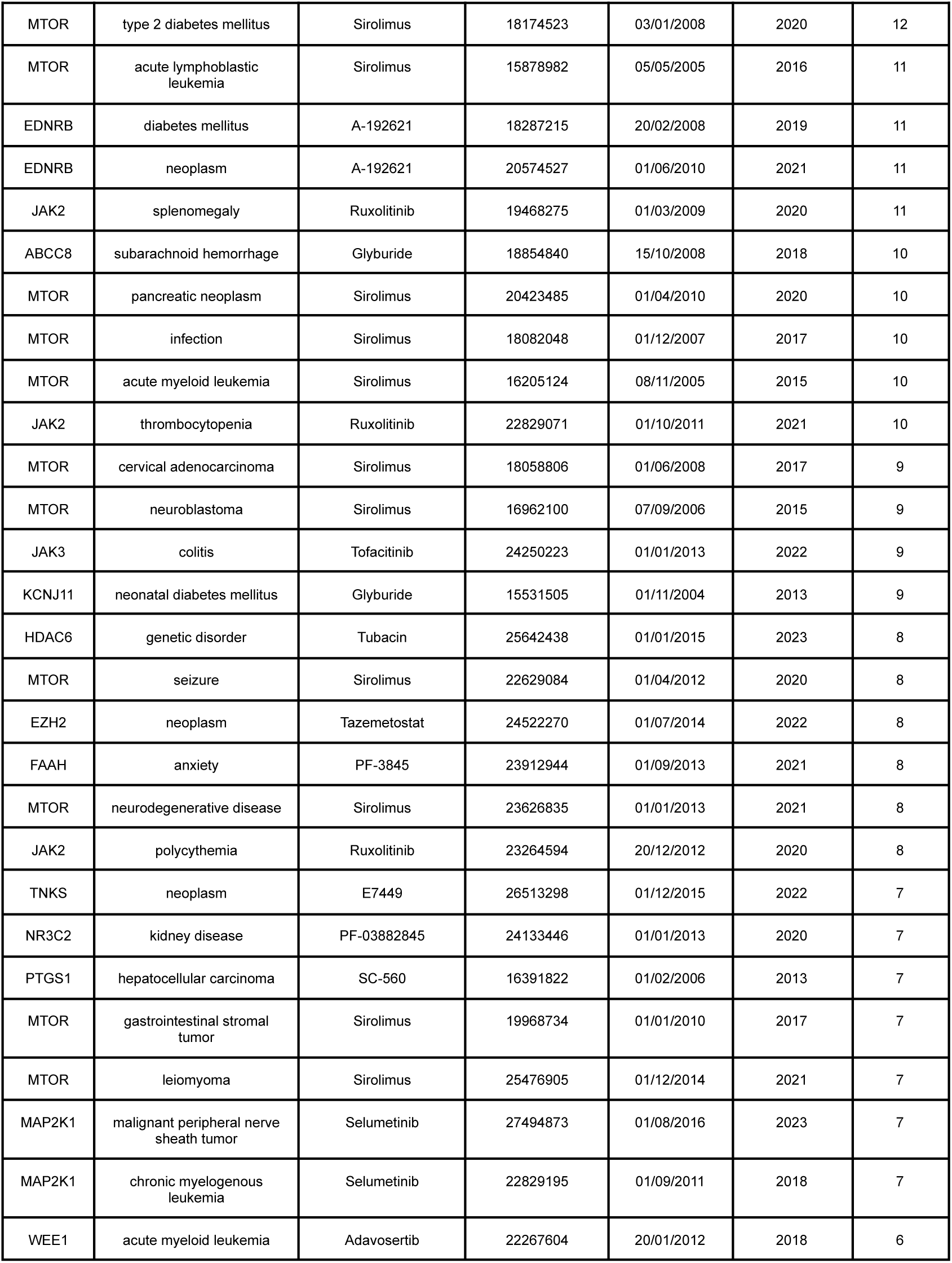
Extreme cases of early T-D evidence from chemical probe studies. Shown are the 30 T-D associations with the largest time lag between an initial chemical probe publication and the first corresponding non-literature evidence in the Open Targets platform. Pairs are ranked by the time span before non-literature evidence to showcase the most prominent examples of chemical probe-based findings preceding other data types. In cases with multiple articles found for the same P-T-D, only the first (largest time lag) was kept in the table.

| Gene Symbol | Disease | Chemical Probe | Article (pmid) | Publication Date | First Evidence Year | Time span (years) |
| --- | --- | --- | --- | --- | --- | --- |
| CCKBR | duodenal ulcer | L-365260 | 9892884 | 01/01/1999 | 2018 | 19 |
| MTOR | rhabdomyosarcoma | Sirolimus | 10029080 | 01/02/1999 | 2013 | 14 |
| MTOR | type 2 diabetes mellitus | Sirolimus | 18174523 | 03/01/2008 | 2020 | 12 |
| MTOR | acute lymphoblastic leukemia | Sirolimus | 15878982 | 05/05/2005 | 2016 | 11 |
| EDNRB | diabetes mellitus | A-192621 | 18287215 | 20/02/2008 | 2019 | 11 |
| EDNRB | neoplasm | A-192621 | 20574527 | 01/06/2010 | 2021 | 11 |
| JAK2 | splenomegaly | Ruxolitinib | 19468275 | 01/03/2009 | 2020 | 11 |
| ABCC8 | subarachnoid hemorrhage | Glyburide | 18854840 | 15/10/2008 | 2018 | 10 |
| MTOR | pancreatic neoplasm | Sirolimus | 20423485 | 01/04/2010 | 2020 | 10 |
| MTOR | infection | Sirolimus | 18082048 | 01/12/2007 | 2017 | 10 |
| MTOR | acute myeloid leukemia | Sirolimus | 16205124 | 08/11/2005 | 2015 | 10 |
| JAK2 | thrombocytopenia | Ruxolitinib | 22829071 | 01/10/2011 | 2021 | 10 |
| MTOR | cervical adenocarcinoma | Sirolimus | 18058806 | 01/06/2008 | 2017 | 9 |
| MTOR | neuroblastoma | Sirolimus | 16962100 | 07/09/2006 | 2015 | 9 |
| JAK3 | colitis | Tofacitinib | 24250223 | 01/01/2013 | 2022 | 9 |
| KCNJ11 | neonatal diabetes mellitus | Glyburide | 15531505 | 01/11/2004 | 2013 | 9 |
| HDAC6 | genetic disorder | Tubacin | 25642438 | 01/01/2015 | 2023 | 8 |
| MTOR | seizure | Sirolimus | 22629084 | 01/04/2012 | 2020 | 8 |
| EZH2 | neoplasm | Tazemetostat | 24522270 | 01/07/2014 | 2022 | 8 |
| FAAH | anxiety | PF-3845 | 23912944 | 01/09/2013 | 2021 | 8 |
| MTOR | neurodegenerative disease | Sirolimus | 23626835 | 01/01/2013 | 2021 | 8 |
| JAK2 | polycythemia | Ruxolitinib | 23264594 | 20/12/2012 | 2020 | 8 |
| TNKS | neoplasm | E7449 | 26513298 | 01/12/2015 | 2022 | 7 |
| NR3C2 | kidney disease | PF-03882845 | 24133446 | 01/01/2013 | 2020 | 7 |
| PTGS1 | hepatocellular carcinoma | SC-560 | 16391822 | 01/02/2006 | 2013 | 7 |
| MTOR | gastrointestinal stromal tumor | Sirolimus | 19968734 | 01/01/2010 | 2017 | 7 |
| MTOR | leiomyoma | Sirolimus | 25476905 | 01/12/2014 | 2021 | 7 |
| MAP2K1 | malignant peripheral nerve sheath tumor | Selumetinib | 27494873 | 01/08/2016 | 2023 | 7 |
| MAP2K1 | chronic myelogenous leukemia | Selumetinib | 22829195 | 01/09/2011 | 2018 | 7 |
| WEE1 | acute myeloid leukemia | Adavosertib | 22267604 | 20/01/2012 | 2018 | 6 |

The co-occurrence of targets and diseases within the chemical probe literature reveals a landscape of focused research priorities. Figure 5 presents a heatmap summarising the associations between the 30 most frequent targets and the 30 most frequent diseases occurring in our dataset, illustrating three key dimensions of evidence: the volume of research (number of unique articles), indicated by colour intensity; the translational status of the pair, represented by the maximum clinical phase reached by any drug for that specific indication (number inside the bubble); and the number of reported chemical tool compounds per T-D pair (number of unique chemical probes), represented by bubble size.

**Figure 5.**
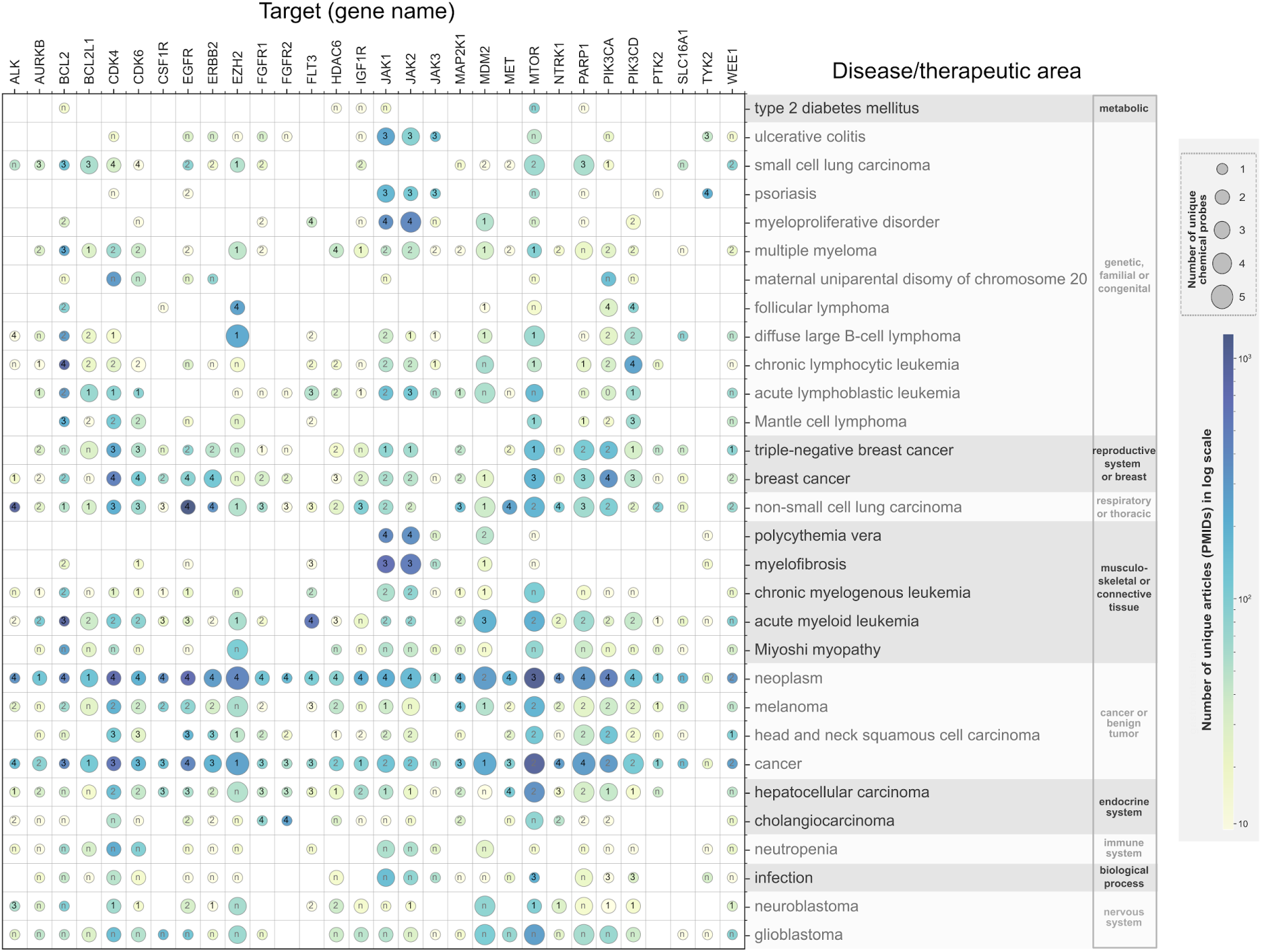
Landscape of T-D links investigated with chemical probes literature for the most frequent targets and diseases. The heatmap illustrates the T-D associations between the 30 most frequent biological targets (abscissa; official gene symbols are given) and the 30 most frequent diseases (ordinate; EFO disease names are given), as well as their adjacent combinations. Data is derived from publications where chemical probes were used as tools to investigate the specific T-D link. Colour intensity corresponds to the number of supporting articles (>9). The size of the bubbles corresponds to the number of different chemical probes used in the respective articles. The number in the bubble reflects the highest clinical phase of drugs (if any available) for the T-D pair (1…phase 1; 2…phase 2; 3…phase 3; 4…approved; n…preclinical). Diseases are grouped by therapeutic area.

The heatmap (Figure 5) visually confirms the observed heavy research concentration in oncology, a trend that transcends specific therapeutic categories. Malignancies constitute the majority of entries not only in the "cancer or benign tumor" section but also across the "respiratory" (e.g., "non-small cell lung carcinoma"), "reproductive" (e.g., "breast cancer"), and "genetic, familial or congenital" groups, which encompass various leukemias and lymphomas. Our dataset reveals that "neoplasm", "cancer", and "non-small cell lung carcinoma" have the largest number of distinct targets, 106, 97, and 58 respectively. For these diseases, there are multiple approved drugs available targeting a variety of different targets (as seen in Figure 5). Nevertheless, compounds in clinical development are still being investigated in the context of alternative targets, such as MDM2, WEE1, and BCL2L1 (for all three cancer types mentioned above). These figures underscore the massive footprint of oncology in the chemical probe literature, likely driven by the routine use of broad selectivity profiling and kinase screening panels.^15^

The plot also enables interpretation of specific T-D associations. For example, the associations between JAK2 and "myeloproliferative disorder", "polycythemia vera" and "myelofibrosis" stand out with dark colouring (corresponding to a high number of associated articles) and large bubbles (corresponding to a high number of chemical probes) containing the number "4" or "3" (approved drug available or clinical candidate in Phase 3). This indicates a relationship that is not only well-documented in the literature but has successfully translated into approved therapies. Similarly, the link between EGFR and "non-small cell lung carcinoma" is prominent, reflecting its status as a clinically validated therapeutic treatment. In contrast, T-D pairs with no clinical annotation represent relationships where chemical probe evidence precedes clinical success. Examples include T-D pairs for the target SLC16A1, which has been investigated in the context of a large number of cancer-related diseases such as "small cell lung carcinoma" and "multiple myeloma" as well as "Miyoshi myopathy" (not cancer-related).

Examples of T-D pairs in which chemical probe evidence precedes drug evidence are of particular interest, as they may signal promising new treatment concepts in drug discovery. However, because the data analyses capture only a static accumulation of P–T–D links, they may also reflect investigations that have not progressed to clinical trials for various reasons. To reliably identify truly novel preclinical therapeutic hypotheses, it is therefore essential to incorporate the temporal dimension.

### Identification of novel preclinical therapeutic opportunities

To identify the next wave of scientific discovery, we extended our analysis to uncover novel P-T-D connections that are currently in the preclinical stage. We applied strict filtering criteria to ensure these associations represent genuine novelty and robustness: selected pairs must be supported by recent evidence (first chemical probe publication <5 years ago), be relatively understudied (≤20 articles) and must have been tested with multiple tool compounds (≥2 distinct chemical probes), yet lack any approved drugs or active clinical programs. The resulting dataset (Supplementary File “ner_probes_triplets_novel.tsv”) contains 135 T-D links shown in Figure 6 which reveals distinct areas of research activity. Supplementary Figure S3 shows an expanded view covering a time span of ten years.

**Figure 6.**
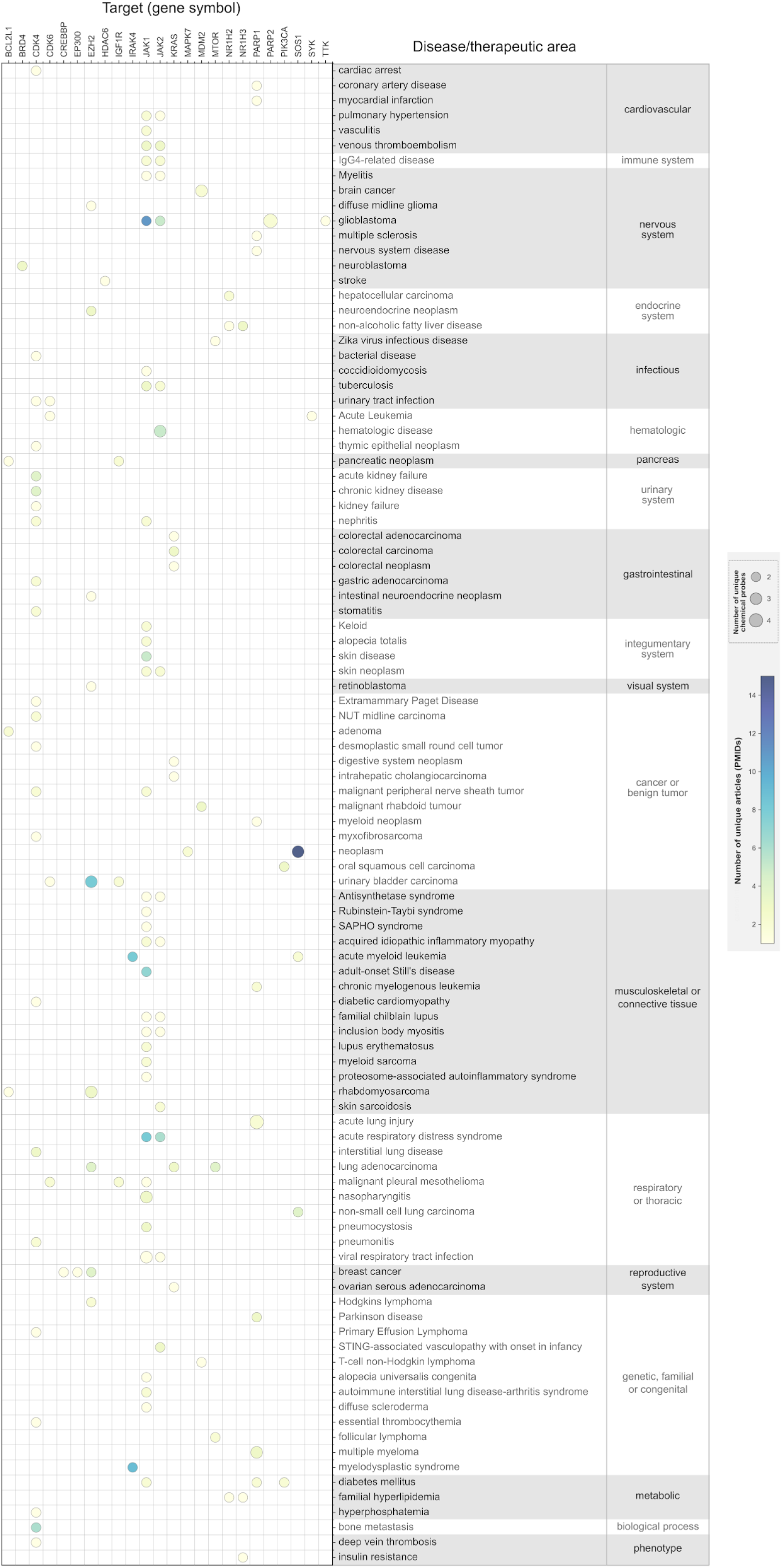
T-D links with novel chemical probe evidence. The heat map illustrates high-confidence, emerging T-D associations derived from chemical probe literature. Pairs were selected based on strict criteria to ensure novelty and robustness: (1) Recent evidence (first chemical probe publication <5 years ago); (2) Robustness (investigated with ≥2 distinct chemical probes); (3) Early-stage volume (<20 supporting articles); and (4) Novelty (no drugs approved or in clinical phases for the pair). Diseases (y-axis) are grouped by Open Targets therapeutic area, and targets (x-axis) are listed by gene symbol. Colour intensity corresponds to the number of supporting articles, while the size of the bubbles corresponds to the number of different chemical probes used.

### Therapeutic repurposing in non-oncology indications

The analysis revealed a group of established oncology and metabolic targets investigated in novel chronic disease contexts (Figure 6). A significant subset of these associations suggests the repurposing of cell cycle and DNA repair modulators for renal and neurological conditions. Specifically, CDK4 was identified in association with acute kidney failure and chronic kidney disease, indicating preclinical investigation into cell cycle regulation as a strategy for renal protection.^16–20^ Similarly, PARP1 was linked to neurodegenerative and inflammatory conditions, including Parkinson’s disease and multiple sclerosis, reflecting literature exploring the inhibition of parthanatos in the central nervous system.^21–23^ In the endocrine and metabolic domain, the nuclear receptor NR1H3 (LXR) displayed associations with non-alcoholic fatty liver disease, familial hyperlipidemia, and insulin resistance, consistent with recent efforts to target LXR signalling in metabolic pathology.^24–26^

### Precision approaches in rare and refractory autoimmunity

A prominent grouping of associations involved the application of Janus kinase (JAK) inhibitors in rare or refractory inflammatory phenotypes (Figure 6). Our pipeline identified JAK1 and JAK2 in the context of specific interferonopathies and autoinflammatory disorders, including COPA syndrome (autoimmune interstitial lung disease-arthritis) for JAK1 and STING disease related to JAK2.^27,28^ Furthermore, Janus kinases have been associated with complex autoimmune conditions such as IgG4-related disease and antisynthetase syndrome for both JAK1 and JAK2, and adult-onset Still’s disease for JAK1.^29–38^ Notably, many of these retrieved associations, particularly COPA syndrome and Still’s disease, highlight a recurring link between JAK modulation and refractory musculoskeletal manifestations. This aligns with the well-established efficacy of JAK inhibitors in common rheumatic diseases, extending their relevance to these rare, complex phenotypes. The analysis also retrieved links between JAK1 and severe dermatological conditions, specifically alopecia totalis and alopecia universalis congenita, aligning with the expanding scope of kinase inhibition in diverse immune-mediated pathologies.^39,40^

#### Targeted strategies in defined oncogenic contexts

Within the oncology domain, the pipeline identified pairs linking specific molecular vulnerabilities to aggressive or hard-to-treat malignancies (Figure 6). MDM2 appeared with a strong signal in the context of malignant rhabdoid tumours, T-cell non-Hodgkin lymphoma and brain cancer, corresponding to research focusing on p53 restoration in specific genetic backgrounds.^41–46^ For genetically defined cancers, the analysis identified EZH2 in association with diffuse midline glioma and retinoblastoma, as well as SOS1 linked to acute myeloid leukaemia.^47–50^ Additionally, IGF1R has been linked to malignant pleural mesothelioma, reflecting the continued search for actionable signalling drivers in this notoriously resistant malignancy.^51,52^

### Chemical probes in the scientific literature and existing target-disease evidence in the Open Targets Platform

To systematically assess the unique value and contribution of chemical probe-derived T-D evidence, we quantitatively investigated the intersection of chemical probe-derived evidence (5,558 unique T-D pairs) with existing T-D association data in the Open Targets Platform (https://platform.opentargets.org/)(Supplementary File “ner_probes_triplets_ptpairs_evd.tsv”). Figure 7A demonstrates the distribution of T-D pairs, identified in the chemical probe literature, across various established evidence types in the platform. Acknowledging that different evidence sources provide complementary information and that their definitive nature often depends on the particular circumstances of the study, our analysis highlights two key roles for chemical probes. In 43.7% of cases, they corroborate associations already supported by other evidence sources such as "somatic mutations", "known drugs", or "genetic associations". Crucially, they also provide significant value by adding functional weight to hypotheses previously reliant on evidence types such as "RNA expression", "animal models", or "literature", thereby serving as a powerful orthogonal validation tool.

**Figure 7.**
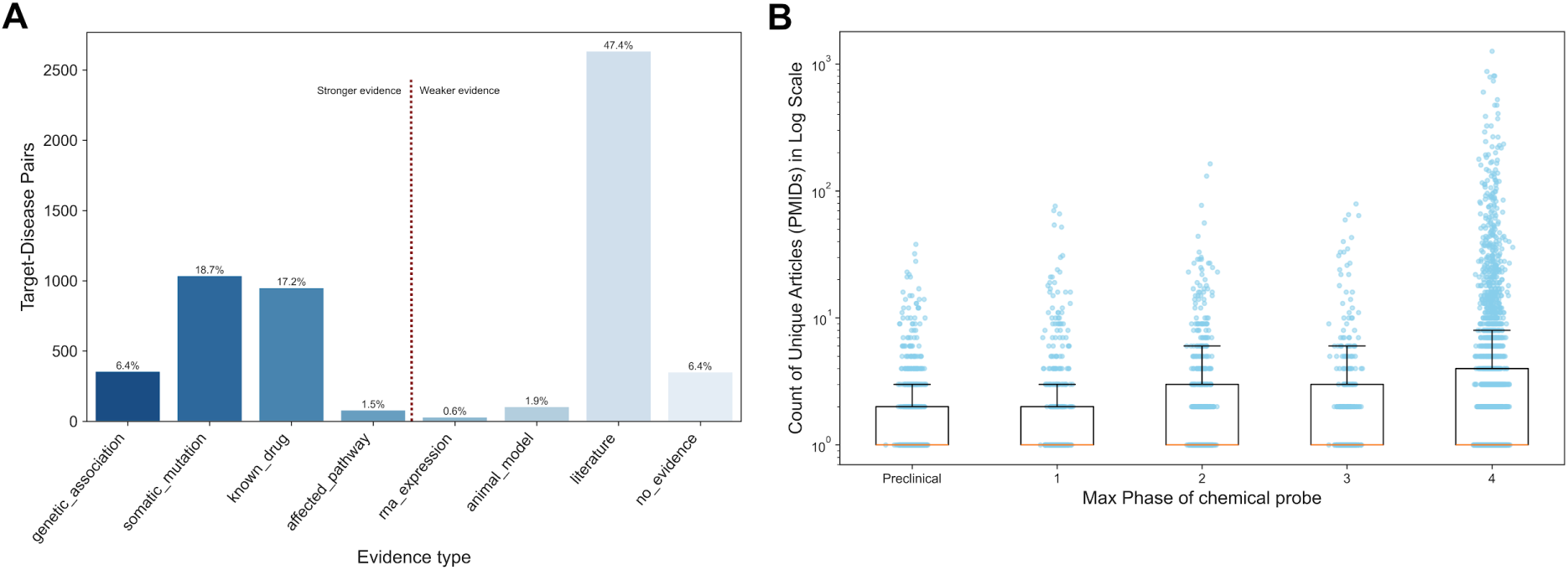
T-D pairs in chemical probe articles supporting existing T-D evidence in the Open Targets Platform and chemical probes evidence from papers from a drug discovery perspective. (A) Distribution of T-D pairs categorised by the type of evidence supporting the association. A vertical dashed line separates evidence types into two categories: ’stronger’ and ’weaker’ evidence types.^10,53^ The value above each bar represents the percentage of total T-D pairs for that specific evidence type. (B) Box plots showing the distribution of the number of unique PubMed IDs (PMIDs) for papers reporting P-T-D triples associated with preclinical and clinical (phases 1-4) drug development stages. The ordinate is on a logarithmic scale. Each dot represents the number of PMIDs for a specific T-D pair within each development phase.

Specifically, papers reporting P-T-D triples provide crucial functional validation and mechanistic insight for numerous T-D pairs primarily supported by evidence categories classified as “weaker”, such as “RNA expression” data (n=33) and "animal models" (n=106). Furthermore, the large number of pairs categorised under "literature" (n=2,636; 47.4% of all pieces of evidence), which represents T-D evidence extracted from the scientific literature using an independent pipeline, suggests that chemical probe experiments often provide direct functional/mechanistic evidence for a T-D link. This can be interpreted as strengthening the evidence available from bioactivity data of (mostly preclinical) compounds reported in these papers.

The identification of T-D 353 pairs (6.4%) where no prior evidence was catalogued highlights the truly discovery-oriented nature of chemical probes. As an example of preclinical studies providing foundational evidence for new hypotheses, a 2020 publication linked the nuclear receptor NR1H2 (LXRβ) to insulin resistance using two distinct LXR agonist chemical probes, GW3965 and T0901317, to successfully restore glucose uptake in a cellular model.^54^ The consistent effect observed with two different chemical probes provides strong functional evidence for this emerging T-D association, representing an opportunity to enrich the existing Open Targets Platform knowledge base.

The Chemical probe literature also captures novel applications for clinical-stage molecules, generating evidence that complements primary indication-focused data. A compelling example involves the target AURKB and osteoarthritis; a study published in 2023 demonstrated that Barasertib, a potent AURKB inhibitor in clinical development for leukaemia, alleviates osteoarthritis in a rat model.^55^ This highlights how a compound developed for oncology can serve as a chemical probe to uncover therapeutic hypotheses in entirely different disease areas.

In addition to therapeutic links, this approach may also be able to identify potential safety liabilities, a type of evidence often structured differently from primary disease associations. For example, a 2020 publication discusses the association between the target FFAR1 and the adverse event hyperbilirubinemia in the context of Fasiglifam, a FFAR1 agonist whose development for diabetes was terminated in Phase 3 clinical trials.^56^ This demonstrates how mining the literature can serve as an early warning system for potential target-mediated toxicities. Finally, this approach can identify high-confidence, clinically relevant links that are lagging in the conventional curation pipeline, thereby expanding coverage with automated extraction. A key example is the association between the epigenetic target EZH2 and extraskeletal myxoid chondrosarcoma. Multiple publications cite clinical trial results (e.g., NCT02601937) for the approved EZH2 inhibitor Tazemetostat, which showed promising anti-tumour activity in patients with this rare cancer.^57,58^

Figure 7B shows the relationship between the volume of supporting literature evidence and the clinical developmental stage of therapeutic hypotheses for all P-T-D triples in the dataset. The analysis reveals a distinct positive correlation: as T-D concepts progress from preclinical stages through clinical Phases 1 to 4, there is a corresponding increase in the number of unique publications associated with them. The logarithmic scale underscores the substantial body of evidence that accumulates for targets advancing successfully through the pipeline.

### Timeline of reporting chemical probes in literature

To investigate the temporal dynamics surrounding chemical probe availability and their subsequent utilization in generating T-D evidence, we analyzed the timelines of chemical probe approval. This is the process that certifies that a molecule meets strict standards for potency, selectivity, and cellular activity, ensuring it is a reliable, high-quality tool for linking a specific protein target to a biological phenotype without misleading off-target effects.^1,4,8,9,59^ Our analysis reveals two distinct phases of development rather than a continuous acceleration (Figure 8A). Following a baseline of limited activity prior to 1998, a distinct inflection point occurs around 2006, marking a step-change in annual chemical probe output. However, the period from 2008 to 2020 is characterized by a stable "steady-state" of discovery, where the annual frequency plateaus rather than continues to rise. This is evidenced by the cumulative count, which follows a linear rather than exponential trajectory during this period, indicating that while the total repertoire of chemical probes is growing, the rate of new chemical probe validation has remained consistent over the last decade.

**Figure 8:**
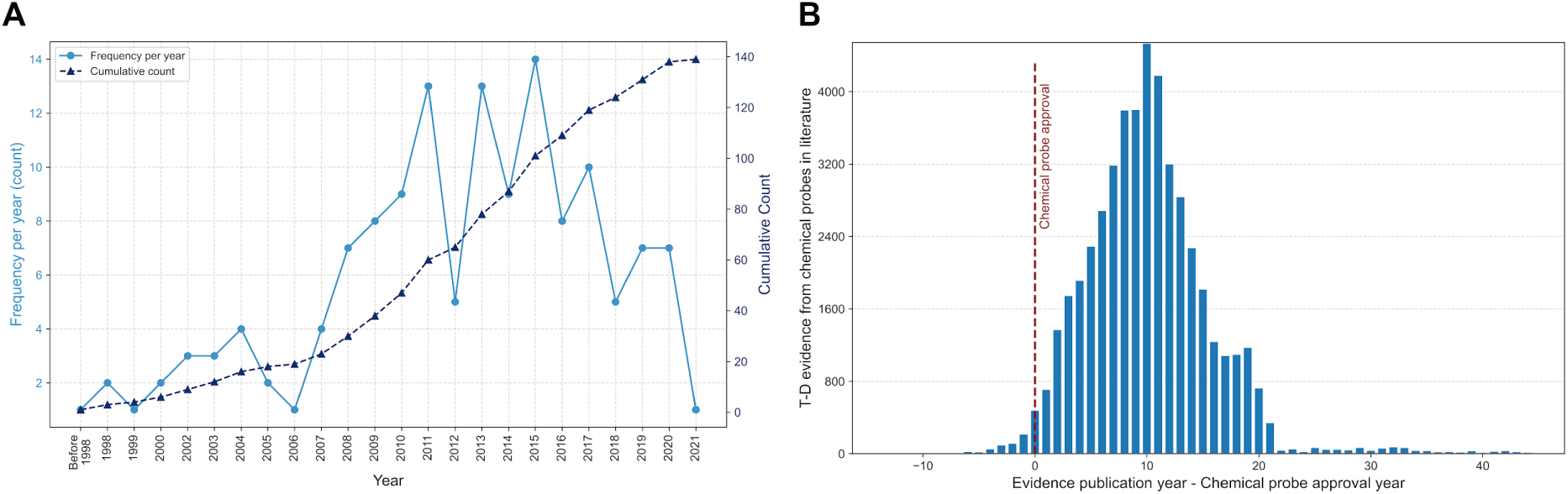
Temporal trends in chemical probe approval and associated T-D evidence publication. (A) Annual frequency (light blue line, left axis) and cumulative count (dark blue dashed line, right axis) of chemical probe approvals from approximately 1988 to 2022, illustrating the increasing availability of characterized chemical probes over time. (B) Distribution histogram showing the time lag (in years) between the year of a chemical probe’s designated approval (Chemical Probe Approval Year) and the publication year of literature evidence supporting a specific T-D association (Evidence Publication Year) using that chemical probe. The x-axis represents the difference (Evidence Publication Year - Chemical Probe Approval Year). The red dashed line indicates the median time lag, highlighting that evidence from the literature often emerges several years after a chemical probe becomes available.

We further examined the time interval between the designated approval year of a chemical probe and the publication year of studies employing that specific chemical probe to provide evidence for a T-D association. The resulting distribution analysis (Figure 8B) reveals a significant positive time lag, characterised by a unimodal distribution with a distinct positive skew (right-skewed). This shape indicates that while a peak frequency of evidence publication occurs roughly 5 to 15 years post-chemical probe approval (confirmed by the positive median lag, red dashed line), a long tail exists where some evidence emerges at considerably later times.

This finding highlights an inherent delay between the introduction of a validated chemical probe and its widespread adoption and application, leading to published T-D linkage evidence within the scientific literature. Extrapolating from these trends, the continued acceleration in chemical probe approvals observed in Figure 8A suggests a rapidly expanding repertoire of chemical probes becoming available for future research. Although the characteristic multi-year time lag for evidence generation depicted by the positively skewed distribution in Figure 8B is likely to persist for the uptake and reporting on *newly* developed chemical probes, the overall volume and rate of published T-D evidence derived from the growing cumulative number of chemical probes is anticipated to increase substantially in the coming years.

### Chemical probe articles as early evidence for target-disease links

Building upon this established role of chemical probes in adding crucial weight, particularly to nascent or weakly supported hypotheses, the investigation next sought to address the temporal dimension of this contribution. Specifically, the analysis aimed to quantify the extent to which chemical probe articles can provide the *first* significant line of evidence for a T-D link, preceding other forms of validation or supporting data.

To determine if chemical probe studies provide early evidence for novel T-D associations, we performed a multi-step temporal analysis. We began with the initial set of 5,558 unique T-D pairs identified from the literature. First, we filtered for pairs that had any dated evidence within the Open Targets Platform, which narrowed our set to 4,558 pairs. To enable a direct comparison between evidence types, this set was further restricted to pairs containing dated non-literature evidence, resulting in a cohort of 1,688 pairs. For this final group, we compared the publication date of the earliest chemical probe article against the date of the earliest non-literature evidence. This analysis identified 185 distinct T-D pairs where the chemical probe publication predated any other catalogued non-literature data. These 185 pairs correspond to 730 specific P-T-D-publication instances, representing cases where chemical probe studies provided the first available evidence for a particular target-disease.

To evaluate this potential for chemical probes to provide early evidence, associations were specifically analysed where the first chemical probe-based evidence publication preceded other evidence types by at least one year (Figure 9). This focused analysis quantifies the typical time lead provided by chemical probes in these instances. When compared against major evidence categories (Figure 9A), the initial chemical probe evidence typically predates the appearance of evidence such as genetic associations, known drug data, or pathway annotation by approximately 1 to 7 years.

**Figure 9:**
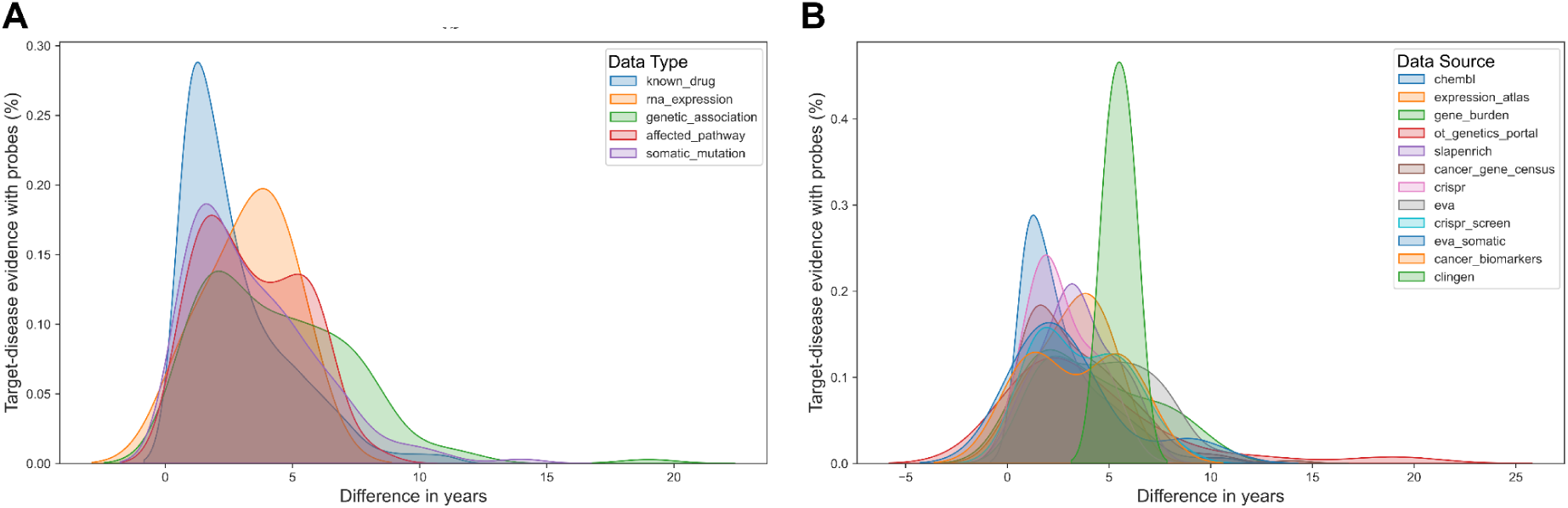
Distribution of time lead for chemical probe evidence preceding other evidence types. These density plots illustrate the distribution of time leads for T-D associations where the first chemical probe evidence was published at least one year before the first appearance of other specified evidence types. The abscissae represents this lead time in years, calculated as First Other Evidence Type Publication Year - First Chemical Probe Evidence Publication Year. (A) Distribution of the lead times (in years) observed when comparing the first chemical probe evidence publication year against the subsequent first appearance year of major evidence categories (“known_drug”, “RNA expression”, “genetic association”, “affected pathway”, “somatic mutation:). (B) Distribution of the lead times (in years) observed when comparing the first chemical probe evidence publication year against the subsequent first appearance year of various granular evidence sources (e.g., “ChEMBL”, Expression Atlas, genetics portals, functional genomics screens, curated cancer datasets) for this filtered dataset.

A more granular comparison against diverse evidence sources (Figure 9B) reveals varying lead times; for example, chemical probe evidence often precedes data from functional genomics screens or other chemical datasets by a relatively shorter period compared to evidence curated in clinical genetics databases (e.g., ClinGen) or comprehensive cancer gene lists, which tend to emerge significantly later.

Among the 185 associations where chemical probe evidence preceded other dated, non-literature evidence in Open Targets, 119 had a lead time of 1-4 years, 54 had a lead time of 5-9 years, and 12 exhibited a significant lead time of 10 years or more, with some approaching two decades.

The most significant gap observed involved the link between CCKBR and Duodenal ulcer, where the initial chemical probe evidence publication(using L-365260, published in 1999)^60^ preceded the first corresponding genetic association data in the Open Targets Genetics Portal (first recorded in 2018) by 19 years. Another notable example is the link between MTOR and Rhabdomyosarcoma, where the initial publication of chemical probe evidence (using Sirolimus/Rapamycin, also published in 1999) preceded the somatic mutation data curated in the Cancer Gene Census (first recorded in 2013) by 14 years. This case highlights how functional evidence derived from a known chemical probe modulating a key pathway component could significantly predate the cataloguing of genetic evidence for the same disease association in established resources. A further instance with a substantial temporal gap was observed for MTOR associated with type 2 diabetes, where chemical probe evidence (using Sirolimus, published 2008)^61^ showed a lead time of 12 years before the appearance of genetic association data in the Open Targets Genetics Portal (first recorded in 2020).

Notably, among these examples is the case involving the chemical probe Sirolimus and its target, MTOR. The findings highlight that studies utilising this single chemical probe provided early evidence linking MTOR function to two phenotypically distinct diseases: rhabdomyosarcoma (a 14-year lead over somatic mutation data) and type 2 diabetes (a 12-year lead over genetic association data).

## Discussion

### Unlocking the early discovery potential of chemical probes

To our knowledge, this study represents the first systematic, large-scale investigation of the chemical probe literature. Our analysis yields four principal findings that collectively redefine the utility of these tools in the drug discovery ecosystem. First, by screening over 260,000 articles using a high-quality dictionary of 561 chemical probes, we successfully mapped the functional landscape of the field, extracting 5,558 unique target-disease (T-D) associations. Second, the application of strict quality filters (recent evidence, ≥2 chemical probes) identified 135 high-confidence novel P-T-D triples which uncovered distinct areas of research activity: notably, therapeutic repurposing in non-oncological indications (such as renal, cardiovascular, and neurodegenerative diseases) and mechanism-based approaches in precision oncology. Third, we established the significant temporal precedence and discovery potential of this evidence; chemical probe data often predates structured information in major knowledge bases by 1 to 7 years on average, with our pipeline identifying 353 pairs (6.4%) that completely lacked prior evidence in the Open Targets platform. Fourth, we demonstrated that chemical probes are essential for strengthening T-D evidence; while they corroborate strong links, their greatest value lies in providing functional validation for associations previously supported only by "weaker" evidence categories, such as RNA expression or animal models.

### Robustness through multiplicity

A key insight from our data is that the strength of a T-D association is best defined not by a single publication, but by the convergence of evidence. While text mining inevitably captures noise from single mentions, the most compelling associations in our dataset are those investigated by multiple independent studies using structurally distinct chemical probes. Reliance on a single chemical probe carries the inherent risk that observed phenotypes stem from off-target interactions rather than the intended target inhibition. However, when different chemotypes targeting the same protein yield consistent phenotypic outcomes, it significantly mitigates the risk that the observed effect is due to off-target toxicity. This cross-verification, or "triangulation" of evidence, provides a level of confidence that arguably surpasses single-line genetic associations, highlighting the unique value of chemical probe-based data in de-risking early-stage targets.^1–3,62,63^

### Mechanism-based safety and toxicity signals

Our analysis reveals that a T-D link in the literature does not always imply therapeutic intent. A distinct subset of the identified associations corresponds to mechanism-based toxicity or safety signals. For instance, the strong association identified between PIK3CA and diabetes mellitus captures the well-documented insulin resistance resulting from on-target PI3K inhibition, rather than a potential diabetes treatment. Similarly, links between JAK inhibitors and infectious complications (e.g., tuberculosis reactivation) reflect the immunosuppressive consequences of this drug class. Far from being false positives, these associations demonstrate the pipeline’s high sensitivity in detecting "on-target" phenotypes, establishing it as a valuable resource for identifying potential adverse events early in the drug development process. While identifying these signals is valuable, automatically distinguishing them from therapeutic indications remains a challenge. Recent methodologies utilising fine-tuned clinical language models have shown great promise in accurately categorising these drug-phenotype relationships, with increasing precision in filtering safety signals from unstructured text.^64–66^ By integrating our comprehensive retrieval capabilities with such context-aware workflows, future iterations could automate the distinction between therapeutic efficacy and safety liabilities, thereby creating a more semantically rich evidence base.

### Challenges in literature reporting and automated extraction

A persistent challenge in the field, and an important context for our findings, is the variability in how chemical probe experiments are reported in scientific papers. Ideally, Methods sections should provide comprehensive details, including chemical probe identifiers, concentrations, treatment durations, sources, and references to primary characterization studies,^8^ yet this standard is frequently unmet. A recent study revealed a suboptimal use of chemical probes in cell-based biomedical research, where only 4% of analysed eligible publications used chemical probes within the recommended concentration range and included inactive compounds as well as orthogonal chemical probes.^67^ In general, many publications offer only cursory mentions of the chemical probe used, lacking the detail necessary for readers to evaluate the appropriateness of the tool or the validity of the experimental design. This issue is often compounded by inadequate descriptions of essential controls, such as inactive enantiomers, and a lack of transparency in the chemical probe selection process itself. The main concern is that researchers may select chemical probes based on historical usage or availability rather than on rigorous, up-to-date characterization data, potentially perpetuating the use of tool compounds with known liabilities without adequate acknowledgement.

Beyond reporting standards, technical challenges in entity recognition, particularly the disambiguation of short, polysemous acronyms, led to specific instances of misannotation. For example, common acronyms like "AA" (Alopecia Areata), "MM" (Multiple Myeloma), and "STS" (Soft Tissue Sarcoma) were occasionally mapped to rare genetic conditions (Aarskog-Scott syndrome, Miyoshi myopathy, and Telomere syndrome, respectively) due to the model prioritising exact dictionary matching over semantic probability. Future iterations of this pipeline will require enhanced disambiguation layers to address these linguistic ambiguities.

Equally important, distinct data loss occurs during the Open Targets entity grounding process,^68^ particularly regarding chemical identifiers, where the extracted chemical mentions are mapped to standardized ChEMBL identifiers.^14^ Consequently, any chemical probe cited in the literature that cannot be strictly resolved to a valid ChEMBL ID, due to synonym mismatches, is systematically excluded from the final dataset. An example of this attrition is the chemical probe JQ1, the archetypal chemical probe for BET bromodomain (BRD) targets. Despite its widespread usage and foundational status in the field, JQ1 was excluded from this iteration of the dataset due to the lack of a valid name-to-ID assignment in the specific ChEMBL lookup utilized at the time of processing. As a result, a substantial volume of high-quality functional evidence linking BRD targets to disease phenotypes was inadvertently omitted. This case underscores that while ontology mapping is a prerequisite for generating structured data, it remains a persistent and non-trivial challenge. Ensuring comprehensive coverage in future large-scale analyses will require more robust fuzzy-matching algorithms and tighter synchronization between text-mining dictionaries and reference chemical databases to prevent the loss of such high-impact tools.

### Defining novelty: from genetic hypothesis to functional reality

When interpreting the temporal patterns and "novelty" of our findings, it is important to consider the structural characteristics of the literature mining approach. Our pipeline specifically identifies articles where a chemical probe, target, and disease co-occur within a single sentence. We observed that a noticeable proportion (38.75%) of this evidence originates from review articles rather than primary research papers. This is likely because reviews frequently summarise established knowledge by grouping these three entities in close proximity, the precise criterion used by our model. In contrast, primary research articles may describe the chemical probe in the Methods section, the target in the Introduction, and the disease relevance in the Discussion, potentially leading to false negatives in a strict sentence-level analysis. Consequently, a "novel" association identified in a recent review may actually reflect a discovery made years prior in a primary study that the pipeline missed due to sentence structure.

However, these technical limitations do not diminish the biological significance of the identified associations. To ensure the genuine novelty of these findings, our analysis applied a rigorous exclusion filter, systematically removing any associations where an approved drug or clinical candidate already existed. This confirms that the "novel" T-D pairs highlighted in our results, such as the repurposing of CDK4 or PARP1, represent true therapeutic white space rather than established clinical knowledge. Furthermore, even if the precise moment of discovery predates the specific text-mined sentence, the evidence identified here represents the first pharmacological or functional validation of the link. While a biological connection may have been suggested earlier via genetic or expression studies, chemical probes serve as the pivotal tools that transition a target from a theoretical genetic hypothesis to a druggable reality. By bridging this gap, the chemical probe-based studies we identified provide the necessary functional de-risking required to justify the initiation of drug discovery programs.

### Future directions

The systematic quantification of chemical probe evidence enables the creation of new scoring metrics, specifically measuring chemical probe diversity and temporal maturity, to formally assess the strength and novelty of target validation. Beyond prioritization, this landscape analysis offers a strategic roadmap for chemical probe development initiatives, such as those led by the Structural Genomics Consortium (SGC), by mapping "dark" targets that possess strong genetic rationale but lack pharmacological tools. Conversely, for targets that already possess high-quality chemical probes but lack disease associations, this dataset highlights under-explored biological space, suggesting where established tools could be effectively deployed in phenotypic screens to uncover novel therapeutic utility. Finally, to maximize the precision of these insights, future technical iterations must focus on refining the NLP pipeline itself, particularly by enhancing entity disambiguation for short acronyms, broader ontology mapping and moving beyond sentence-level co-occurrence to capture complex experimental details hidden within full-text documents.

## Methods

### Pilot dataset construction and validation

As an initial proof of concept, a pilot dataset was constructed to explore articles mentioning chemical probes and their use to test target-phenotype links. A simple chemical probe dictionary was constructed, utilising the "good quality set" of Probes&Drugs^8^ (excluding the dataset of Chemicalprobes.org) retrieved in July 2022. To maximize recall, this dictionary was augmented with synonyms for each chemical probe and target, derived from both Probes&Drugs portal (https://www.probes-drugs.org/home/) and ChEMBL database (ChEMBL34).^14^

A natural language processing (NLP) pipeline was developed to analyze articles from a dictionary-based annotated dataset provided by Europe PMC.^68^ This dataset annotates "target", "disease", and "chemical" entities within free text up to 2021. The search engine focuses on articles exhibiting the co-occurrence of a chemical probe (chemical entity), its corresponding target (target entity), and a phenotype (disease entity). More precisely, it identified sentences containing the three entities within the same sentence, which were labelled as a strong link. However, instances where the disease label appeared elsewhere within the article, independent of the chemical probe and target co-occurrence sentence, were also kept and labelled as “weak” links. Further filtering steps were carried out: (a) only articles mentioning SGC chemical probes were considered (compound set=28 in Probes&Drugs), (b) excluding articles where the chemical probe was mentioned in the article’s section "Other", and (c) keeping only articles with activity unit keywords mentioned to increase the likelihood of existence of reports of experimental measurements for the chemical probes mentioned in the articles. The filter was constructed using common unit terms (e.g., nM, μM, nMol, etc.) in ChEMBL activities as part of a regular expression to capture such strings in the article.

The subset underwent a two-stage validation process: (1) manual curation by expert curators to ensure accuracy and consistency of the identified P-T-D triples (strong and weak links), and (2) extraction of bioactivity data and all associated metadata (in analogy to ChEMBL’s literature data extraction). This dataset is available as part of ChEMBL 36 release at Src_ID 72.

### Chemical probes main dataset construction

The dataset was constructed using article annotations from EuropePMC (updated in September 2024). These annotations were generated by a novel, AI-based Named Entity Recognition (NER) model, which replaces EuropePMC’s legacy dictionary-based system.^68^ The model identified three key biological entities: chemicals, targets, and diseases. Subsequently, Open Targets processed these raw annotations to normalize (or "ground") each entity to a standard identifier. Specifically, targets were mapped to Ensembl IDs, diseases to Experimental Factor Ontology (EFO), MONDO, or HP terms, and chemicals to ChEMBL IDs.

The complete literature annotated dataset can be found in the Open Targets FTP repository https://ftp.ebi.ac.uk/pub/databases/opentargets/platform/latest/intermediate/literature_match. A dedicated dictionary of high-quality (HQ) chemical probes was constructed to support the text-mining pipeline by aggregating information from established public resources. Specifically, data were compiled from the Structural Genomics Consortium (SGC) chemical probes list (reflecting updates as of April 2022), Open Science Probes (updated as of April 2023), and ChemicalProbes.org (updated as of April 2023). To ensure the inclusion of well-validated tools suitable for biological investigation, specific quality criteria were applied for chemical probes originating from the ChemicalProbes.org dataset. Only those achieving that resource’s rating of more than three stars in cellular assays ("in cell") and more than three stars for in vivo studies were selected for inclusion. This aimed to enrich the final HQ dictionary with chemical probes deemed particularly well-suited and validated for both cell-based and in vivo experimentation according to those specific criteria. Chemical probes’ and targets’ synonyms, together with other features related to the source sets, were obtained from Probes&Drugs portal and ChEMBL database.

Articles were explored for the co-occurrence of the three entities within the same sentence, with a dictionary-based filtering step applied to enhance confidence, retaining only associations where the identified target was an accepted, validated target for the specific chemical probe mentioned. To further refine the dataset, manual curation was performed on the subset of novel T-D associations to identify disease identifiers wrongly assigned by the NER model—predominantly false positives caused by ambiguous abbreviations. Based on these findings, a blacklist of problematic terms (Supplementary File disease_terms_black_list.csv) was constructed and subsequently used to filter the entire dataset, ensuring these erroneous associations were excluded from all downstream analyses.

### Extracting evidence from Open Targets

Existing T-D evidence in Open Targets Platform using the "targetId" and the "diseaseId" keys. Note that multiple evidence can be found for the same target and disease pair. Evidence was categorized according to the "datatypeId" in two groups: strong evidence and weak evidence. The strong evidence compasses all evidence from “genetic mutations”, “known drugs”, “genetic associations”, or “affected pathways”. The weaker group includes all evidence coming from “RNA expression”, “animal models” or “literature”. Cases where no evidence was found in Open Targets were labelled as "No evidence". Open Targets data was obtained from the public FTP repository (v25.09;http://ftp.ebi.ac.uk/pub/databases/opentargets/platform/25.09/output/association_by_datasource_direct)

### Adding the Open Targets time stamp to the T-D evidence in Open Targets

Timestamp data associated with T-D evidence was obtained from an internal OpenTargets database snapshot dated January 16th, 2025. This dataset contained evidence timestamps which can originate from an available curation date, a linked publication date or a study year (Supplementary Figure 4) and it is linked to specific data sources (datasourceId).^69^ To incorporate this temporal information, the timestamp dataset was merged with the core T-D association dataset using a left join operation, matching records based on diseaseId and targetId. This approach preserved all records from the T-D association dataset (the left table in the join), appending timestamp information where available and retaining records even if no corresponding timestamp was found.

### Assigning a clinical phase to chemical probes

The maximum clinical development phase for each chemical probe within the primary dataset was determined using data retrieved from ChEMBL (version 35). For each chemical probe, its assigned ChEMBL ID (CHEMBL_ID) was utilized to query the MAX_PHASE value from the molecule_dictionary table. The specific SQL query structure employed was: SELECT d.MAX_PHASE FROM molecule_dictionary d WHERE d.CHEMBL_ID = "{probe_chembl_id}". The resulting MAX_PHASE values were interpreted according to the ChEMBL database definitions as follows: NULL (Preclinical), -1 (Unknown), 0.5 (Early Phase 1), 1 (Phase 1), 2 (Phase 2), 3 (Phase 3), or 4 (Approved).

### Extracting evidence from Open Targets

Therapeutic area categories were retrieved from the Open Targets Platform. The "therapeuticAreas" column was used to extract all associated parent terms from the disease ontology tree. In cases where a disease was mapped to multiple therapeutic area categories, a primary category was assigned based on a hierarchical prioritisation of the ontology source: Open Targets identifiers were prioritised first, followed by EFO IDs, MONDO IDs, and finally other sources. Open Targets data was obtained from the public FTP repository (v25.09; http://ftp.ebi.ac.uk/pub/databases/opentargets/platform/25.09/output/disease).

### Retrieval of clinical drug evidence

Information regarding approved drugs and clinical candidates was obtained from the Open Targets Platform. We mapped this clinical data to our dataset by matching the unique "targetId" and "diseaseId" identifiers for each target-disease pair. The specific column "phase" was utilised to extract the clinical development stage, enabling the identification of pairs with existing therapeutic interventions. For further details on the specific definitions and classification of these clinical phases, see previous section on ChEMBL chemical phase or refer to the Open Targets documentation (https://platform-docs.opentargets.org/drug/clinical-precedence). Open Targets data was obtained from the public FTP repository (v25.09; http://ftp.ebi.ac.uk/pub/databases/opentargets/platform/25.09/output/known_drug)

### Constructing chemical probes timelines

To study the impact of chemical probes over time, a "Probe Approval Year" was first established for each compound. This was defined as the publication year of the earliest primary article that characterized the compound as a chemical probe. This information was retrieved from the Probes&Drugs portal, which typically links to the original publication from sources such as the ChemicalProbes.org portal. If a chemical probe was linked to multiple primary publications (Supplementary Figure 2), the earliest date was always chosen.

Next, the time lag (in years) between a chemical probe’s approval and its application in a study supporting a specific T-D association was calculated. This was calculated as:

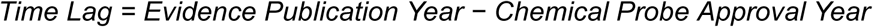

### Calculating the time lag between the chemical probe’s evidence and other evidence

T-D pairs associated with time-stamped evidence (publication year) were selected for this temporal analysis. To mitigate potential redundancy, given that chemical probe evidence derives from literature sources also processed by Open Targets, comparisons were made exclusively against non-literature evidence types obtained from Open Targets. For each qualifying T-D pair, the publication year associated with chemical probe evidence (probe_year) and the earliest year for any time-stamped non-literature evidence (nonLit_year) were identified. A time interval was calculated by subtracting the chemical probe evidence year from the non-literature evidence year (ΔYear = nonLit_year - probe_year).^69^ Subsequently, only T-D pairs exhibiting a positive time interval (ΔYear > 0), indicating that the chemical probe evidence publication preceded the earliest recorded non-literature evidence, were retained for further analysis. Cases with a positive time interval were then grouped based on the datatypeId of the corresponding non-literature evidence (specifically: known_drug, rna_expression, genetic_association, affected_pathway, somatic_mutation) and by the associated datasourceId (including chembl, expression_atlas, gene_burden, ot_genetics_portal, slapenrich, cancer_gene_census, crispr, eva, crispr_screen, eva_somatic, cancer_biomarkers, clingen). The distribution of these time intervals within each category was visualised using density plots.

## Conclusion

In this study, we provide the first systematic, large-scale quantification of chemical probe literature, redefining these tools as primary engines of early therapeutic discovery rather than simple validation reagents. Our analysis yields four principal findings:

1. **Landscape mapping:** By leveraging a high-quality NLP pipeline across more than 260,000 articles, we successfully mapped the functional landscape of the field, extracting 5,558 unique target-disease associations.
2. **Temporal precedence:** We established the significant temporal precedence of this evidence, revealing that chemical probe data frequently predates structured genetic or clinical information in major knowledge bases by 1 to 7 years.
3. **Repurposing opportunities:** The application of strict quality filters uncovered high-confidence novel associations, highlighting distinct opportunities for therapeutic repurposing in non-oncological indications and mechanism-based precision oncology.
4. **Evidence elevation:** We demonstrated that chemical probes are essential for elevating evidence, providing crucial functional validation for associations previously supported only by weaker, correlative data such as RNA expression or animal models.

Future efforts should focus on formally integrating these chemical probe-derived metrics into target prioritisation frameworks and developing context-aware models to automate the extraction of nuanced experimental details currently buried in the scientific literature.

## Data availability

All data and associated code to reproduce the results presented in this manuscript can be found in our public GitHub repository: https://github.com/chembl/chemical_probes_lit. The primary dataset generated and analysed during the current study, together with all intermediate files, are available in the Zenodo repository: 10.5281/zenodo.17085504.

## Acknowledgements

The authors thank Shyama Saha and Santosh Tirunagari (Literature Services Team, EMBL-EBI) for their insightful discussions, to Ctibor Skuta for sharing Probes&Drugs original publications data and to Tevfik Kizilören (Chemical Biology Services team, EMBL-EBI) for his technical support. This work was conceived and funded by Open Targets, under OTAR2047 "Enhanced Molecular and Clinical Data".The authors also acknowledge financial support from the Wellcome Trust [104104/A/14/Z, 218244/Z/19/Z, 228142/Z/23/Z] and from the Member States of the European Molecular Biology Laboratory (EMBL).

## Author’s contribution

M.F.A., A.R.L. and B.Z. conceived and designed the study. M.F.A. developed the methodology, performed the investigation and formal analysis, contributed to data curation, and wrote the manuscript draft. D.O., I.L. and E.M. contributed to the conceptualization and methodology. H.D. contributed to data curation. N.O. contributed to the conceptualization.

A.R. L. and B.Z. supervised the project. All authors contributed to writing of the manuscript, and reviewed and approved the final manuscript

