## Supplementary files and figures for "Chemical Probes in Scientific Literature: Expanding and Validating Target-Disease Evidence"

The primary dataset generated and analysed during the current study, together with all supplementary files, are available in the Zenodo repository: 10.5281/zenodo.17085504.

**pilot_triples.csv:** articles found in the pilot study with an SGC chemical probe, target and disease. The file reports all triples found in articles.
**probes_HQ.csv:** High-quality probes dictionary used in the automated approach.
**ner_all_triplets.tsv:** first step for the systematic approach with all articles annotated with 3 entities [any chemical, target, disease] if they are found within the same sentence.
**ner_probes_triplets_ptpairs.tsv:** main dataset for the systematic approach with all articles annotated with 3 entities [chemical probe, target, disease] if they are found within the same sentence. The found target is one of the approved targets for the specific chemical probe.
**ner_probes_triplets_ptpairs_evd.tsv:** main dataset for the systematic approach with 3 entities and existing T-D evidence from the OpenTarget Platform (OTP) if available. It also includes details on dated evidence if found, OTP therapeutic area for disease entity, and OTP known drugs/clinical phase for T-D pairs.
**ner_probes.tsv:** all chemical probes found in the main dataset with the corresponding ChEMBL id (probeId), the chemical probe name (probeName), the total number of articles (pmids) they appear in, the number of unique diseases they are linked to (diseases), and the number of unique targets (targets). They are sorted by frequency with respect to the pmids.
**ner_targets.tsv:** all targets found in the main dataset with the corresponding Ensembl target id (targetId), the target's gene symbol (geneName), the total number of articles (pmids) they appear in, the number of unique diseases they are linked to (diseases), and the number of unique chemical probes (probes). They are sorted by frequency with respect to the pmids.
**ner_diseases.tsv:** all diseases found in the main dataset with the corresponding ontology disease id (diseasetId), the disease's name (diseaseName), the total number of articles

(pmids) they appear in, the number of unique targets they are linked to (targets), and the number of unique chemical probes (probes). They are sorted by frequency with respect to the pmids.
**ner_target_disease_pairs.tsv:** all unique target-disease (T-D) pairs from the main dataset sorted by frequency with respect to the number of articles they appear in (pmids).
**ner_probes_triples_novel.tsv:** the subset contains the novel T-D pairs following the strict filter criteria: selected pairs must be supported by recent evidence (first probe publication <5 years ago), not overly studied (≤ 20 articles) and must have been tested with multiple tools (≥2 distinct chemical probes), yet lack any approved drugs or active clinical programs.
**disease_terms_black_list:** A compilation of disease identifiers excluded from the analysis. These terms were flagged during manual curation as systematic false positives within the NER dataset, primarily due to ambiguous abbreviations that led to incorrect disease assignments.

### Supplementary figures

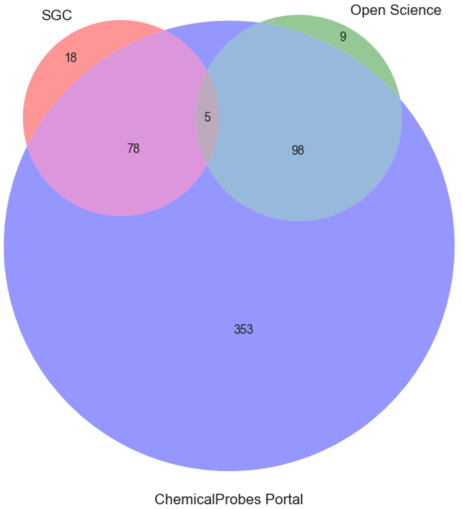


**Supplementary Figure 1. Overlap of High-Quality Chemical Probe Sources.** The Venn diagram illustrates the distribution and overlap of probes from the three curated sources used to build our high-quality chemical probe dictionary: the Structural Genomics Consortium (SGC), Open Science, and the ChemicalProbes.org portal. The numbers in each segment represent the count of unique probes. A total of 561 unique probes were aggregated for this study, with the diagram highlighting the contribution of each source.

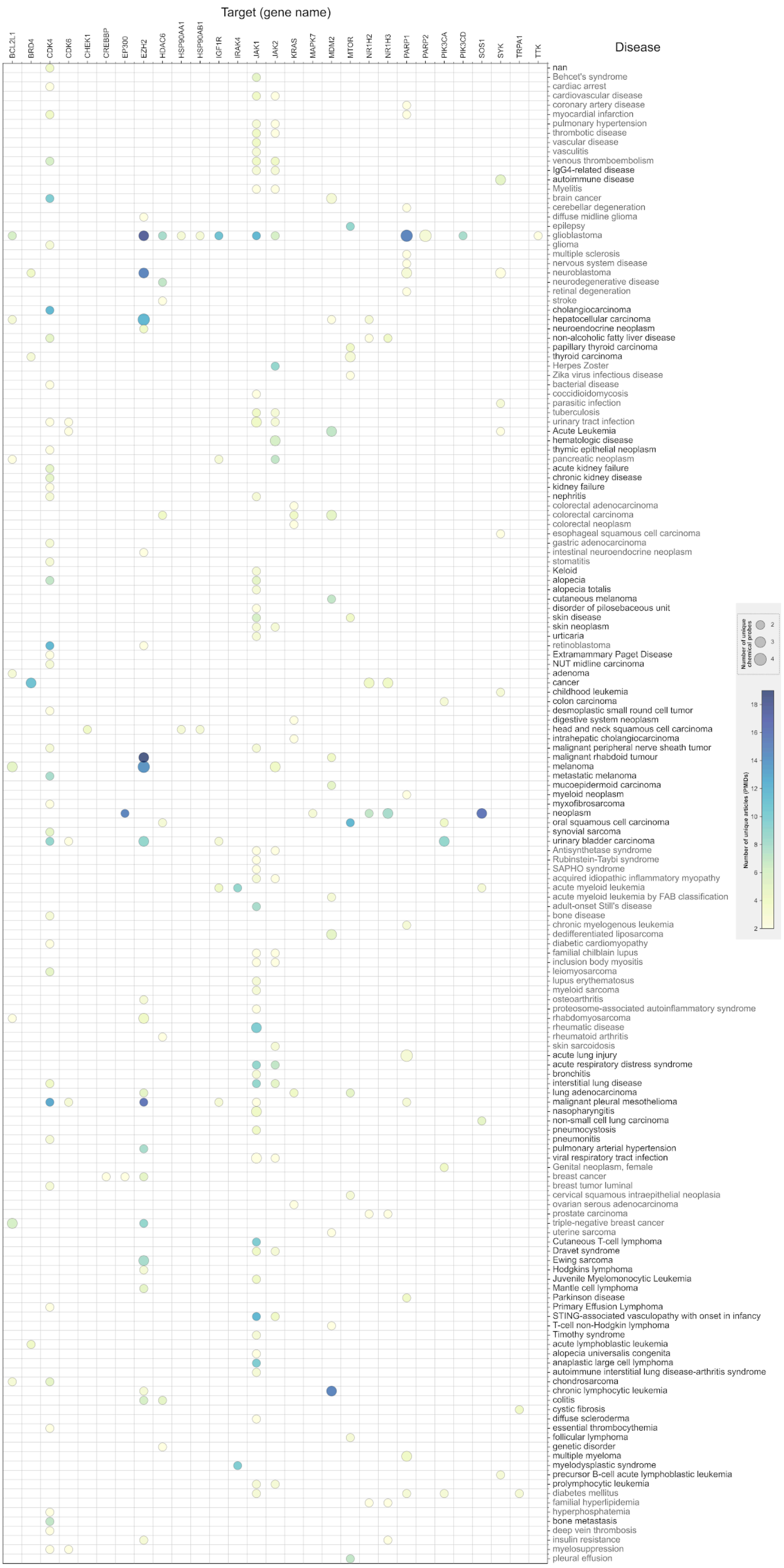


**Supplementary Figure 2. Expanded Novel T-D links with recent chemical probe evidence from the last 10 years.** The heat map illustrates high-confidence, emerging T-D associations derived from chemical probe literature. Pairs were selected based on strict criteria to ensure novelty and robustness: (1) Recent evidence (first chemical probe publication <10 years ago); (2) Robustness (investigated with ≥2 distinct chemical probes); (3) Early-stage volume (2–20 supporting articles); and (4) Novelty (no drugs approved or in clinical phases for the pair). Diseases (y-axis) are grouped by Open Targets therapeutic area, and targets (x-axis) are listed by gene symbol. Colour intensity corresponds to the number of supporting articles, while the size of the bubbles corresponds to the number of different chemical probes used.

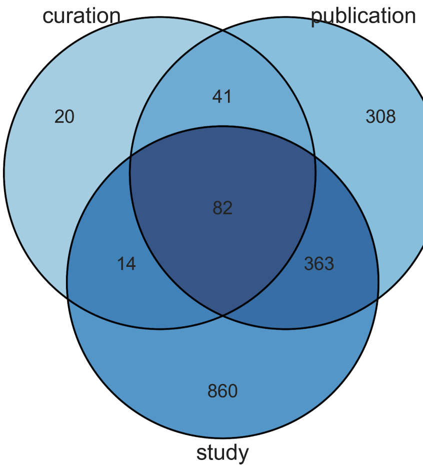


**Supplementary Figure 3. Overlap of Dating Methodologies for Non-Literature Evidence in Open Targets.** This Venn diagram illustrates the distribution of the dating methodologies employed by the Open Targets time stamp pipeline for the 1,688 T-D associations used in our temporal analysis. The three sets represent the main approaches from the platform: a specific curation date, a date from an associated publication, or the start/end date of a clinical study. The diagram shows the degree of overlap, revealing how many associations are dated by single versus multiple methodologies.

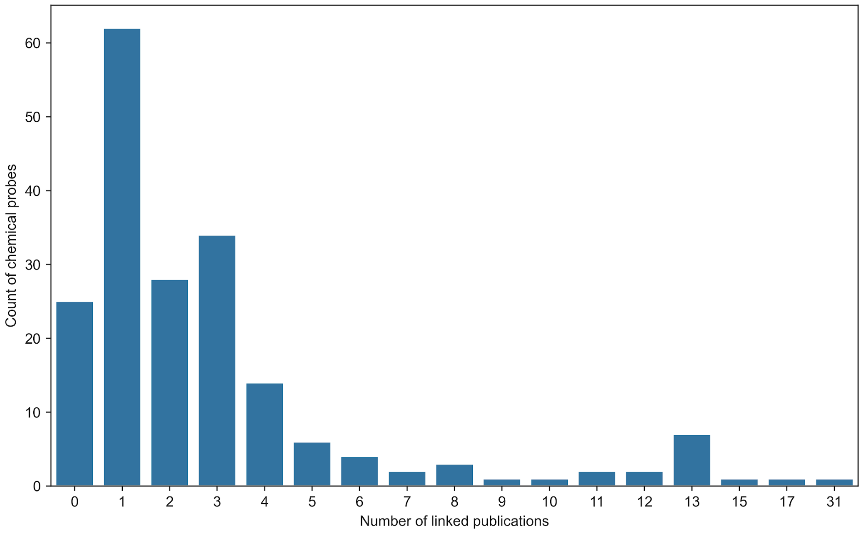


**Supplementary Figure 4. Distribution of Primary Publications Linked to Each Chemical Probe.** This bar chart shows the number of chemical probes (Y-axis) associated with a specific count of primary characterization publications (X-axis), as sourced from the Probes&Drugs portal. The distribution highlights that most chemical probes are linked to a small number of publications (typically 1–3), while very few have a large number of associated articles. This figure provides context for the method used to determine the "Probe Approval Year," where the earliest of these linked publications was selected.
